# Discrete interneuron subsets participate in GluN1/GluN3A excitatory glycine receptor (eGlyR)-mediated regulation of hippocampal network activity throughout development and evolution

**DOI:** 10.1101/2025.09.24.678342

**Authors:** June Hoan Kim, Anna Vlachos, Vivek Mahadevan, Adam P Caccavano, Tue Banke, Oliver C Crawley, Ana I Navarro, Xiaoqing Yuan, Daniel Abebe, Steven Hunt, Geoffrey A Vargish, Ramesh Chittajallu, Mark A G Eldridge, Reza Azadi, Alex C Cummins, Anne-Claire Tangen, Peyton Harmon, Anya Plotnikova, Arya Mohanty, Elisabetta Furlanis, Yating Wang, Min Dai, Brenda Leyva Garcia, Ding Liu, Zongjian Zhu, Hongjie Yuan, Samantha L Summer, Matthew P Epplin, Dennis C Liotta, James Pickel, Bruno B Averbeck, Isabel Perez-Otaño, Jordane Dimidschstein, Gord Fishell, Stephen F Traynelis, Chris J McBain, Kenneth A Pelkey

## Abstract

Decades of studies implicating GluN3A *N*-methyl-D-aspartate receptor (NMDAR) subunits in physiological and pathological function have largely been interpreted through direct regulation of conventional glutamatergic NMDARs. However, emerging evidence indicates that GluN3A frequently assembles with GluN1 forming unconventional glutamate-insensitive NMDARs that operate as native excitatory glycine receptors (eGlyRs). Here we demonstrate that hippocampal somatostatin and neurogliaform interneurons (Sst-INs and NGFCs) express functional eGlyRs from early postnatal through adult ages. In the developing hippocampus eGlyR-mediated excitation of NGFCs with ambient glycine dramatically increases GABAergic tone, with consequences for the generation of giant depolarizing potentials (GDPs). In the mature hippocampus, eGlyR- mediated excitation of Sst-INs regulates sharp wave ripples (SWRs). Finally, we reveal evolutionary conservation of hippocampal Sst-IN eGlyRs and eGlyR- mediated SWR regulation in non-human primates confirming functional eGlyR availability for therapeutic potential in higher species. Our findings underscore that eGlyR mediated regulation of cell and circuit excitability through both cell autonomous and cell non-autonomous mechanisms must be considered to understand GluN3A roles in brain development, plasticity, and disease.

## Introduction

For almost half a century NMDARs have been recognized as crucial mediators of excitatory glutamatergic synapse development, function, and plasticity in brain health and disease. The coincidence detecting features, kinetics, and calcium permeability of conventional GluN1/GluN2 subunit-containing NMDARs confer integrative properties uniquely suited to controlling synaptic maturation, integration, and activity-dependent plasticity [Hansen 2021, Nicoll 2017]. Interestingly, GluN3A NMDAR subunits are also strongly implicated in nascent synapse and circuit development/plasticity [Das 1998, Gonzalez-Gonzalez 2023, Hurley 2024, Sasaki 2002, Tong 2008, Roberts 2009, Henson 2012, Mohamad 2013, Kehoe 2014, Perez-Otano 2016, Crawley 2022]. However, emerging evidence indicates that GluN3A subunits primarily co-assemble with GluN1 to yield unconventional glycine (Gly) gated “non-Hebbian NMDARs” that are glutamate insensitive and voltage-independent with low Ca^2+^ permeability [Chatterton 2002, Pina-Crespo 2010, Awobuluyi 2007, Madry 2007, Kvist 2013a, Bossi 2022]. As such GluN1/GluN3 NMDARs are not members of the glutamate receptor family and are referred to as eGlyRs to differentiate them from both conventional glutamate/glycine gated NMDARs and inhibitory chloride conducting pentameric glycine-gated receptors [Chatterton 2002, Otsu 2019, Crawley 2022, Bossi 2022, Hurley 2024]. Curiously, eGlyRs are often expressed by neurons in forebrain regions essentially devoid of dedicated glycinergic afferents [Zeilhofer 2005, Bossi 2022, Bossi 2023]. Thus, evidence for prototypical rapid, phasic communication through eGlyRs is lacking. Instead, evidence suggests that eGlyRs regulate neuronal excitability by sensing ambient levels of extracellular Gly consistent with observations that surface GluN3A subunits comprise a highly diffusive receptor pool that does not strongly localize to postsynaptic sites [Perez-Otano 2006, Otsu 2019, Bossi 2022, Gonzalez-Gonzalez 2023].

Gly gates eGlyRs by binding to GluN3 subunits, whereas Gly binding to GluN1 profoundly desensitizes eGlyRs [Awobuluyi 2007, Madry 2007, Kvist 2013a, Cummings 2017, Grand 2018, Zhu 2020, Rouzbeh 2023]. While saturating levels of ambient Gly for GluN1 have historically confounded interrogation of eGlyRs, findings that the glycine site antagonist CGP-78608 (CGP) blocks desensitization and awakens eGlyRs has fostered a new era of cellular and network investigation into this notoriously enigmatic receptor subpopulation that has remained elusive since their discovery in recombinant systems over 20 years ago [Chatterton 2002, Matsuda 2002, Yao 2006, Madry 2007, Grand 2018, Otsu 2019, Zhu 2020, Bossi 2022, Rouzbeh 2023, Hurley 2024]. Genetic deletion of *Grin3a*, has revealed critical roles for eGlyRs in regulating diverse behaviors including sensorimotor gating, sleep, place aversion conditioning and fear memory stability [Otsu 2019, Bossi 2022, Brody 2005, Sunagawa 2016, Crawley 2022]. Moreover, mutations or alterations in *GRIN3A/Grin3a* expression in humans and mice are linked to diverse neurological conditions such as schizophrenia, bipolar disorder, addiction, epilepsy, and Huntington’s disease making GluN3A an attractive candidate for therapeutic intervention [Mueller 2004, Gallinat 2007, Marco 2013, Yuan 2013, Yang 2015, Lee 2016, Huang 2017, Marco 2018, Crawley 2022]. The unique features of eGlyR ligand binding, channel activation, and gating can be successfully exploited to develop selective allosteric modulators [Kvist 2013a, Kvist 2013b, Zhu 2020, Rouzbeh 2023, Michalski 2024, Jacobs 2025]. However, determination of therapeutic potential requires thorough characterization of eGlyR function within discrete receptor expressing cell-types across divergent circuits throughout development and across species.

*Grin3a* expression is particularly high in the developing hippocampus but is not ubiquitous exhibiting unique regional and cell type specific patterns with divergent temporal profiles [Sucher 1995, Wong 2002, Roberts 2009, Kehoe 2014, Murillo 2021, Hurley 2024]. At early postnatal stages *Grin3a* is highly expressed in CA1 pyramidal neurons (PNs) throughout the entire rostrocaudal axis then dramatically declines in dorsal CA1 in adults but remains elevated in ventral CA1 pyramidal neurons [Murillo 2021, Hurley 2024]. *Grin3a* is not expressed in developing or adult PNs of CA2, CA3, or dentate gyrus indicating that any potential role for *Grin3a* in regulating circuit maturation and function in these hippocampal substructures requires cell non-autonomous mechanisms [Sucher 1995, Murillo 2021, Beesley 2022, Pizzamiglio 2025]. Hippocampal and neocortical medial ganglionic eminence (MGE)-derived inhibitory GABAergic interneurons express conspicuously high levels of *Grin3a* throughout development, and adult somatostatin- expressing interneurons (Sst-INs) exhibit robust eGlyR-mediated currents [Pfeffer 2013, Paul 2017, Murillo 2021, Bossi 2022, Pizzamiglio 2025]. Individual local circuit interneurons widely innervate large numbers of nearby PNs to critically instruct nascent circuit connectivity through early postnatal depolarizing GABAergic drive [Ben-Ari 2002, Ben-Ari 2007, Cossart 2022, Pelkey 2017]. Moreover, in mature hippocampus GABAergic inhibition from divergent interneuron subtypes imposes discrete spatiotemporal profiles of inhibition to entrain principal cell assemblies and coordinate network oscillations critical for cognition [Klausberger 2008, Stark 2014, Pelkey 2017, Tzilivaki 2023]. These central roles of GABAergic interneurons could be leveraged by ambient Gly through eGlyRs to control network excitability for circuit maturation and rhythmogenesis. Here using a multiparametric approach including transcriptional profiling, conditional loss of function mice, immunohistochemistry, and *ex vivo* electrophysiology with optogenetics and pharmacology we demonstrate that hippocampal Sst-INs and neurogliaform interneurons (NGFCs) express functional eGlyRs from early postnatal through adult ages. Activation of eGlyRs on these interneuron subsets has the capacity to regulate the dominant spontaneous network pacing activities, GDPs and SWRs, in developing and mature hippocampus respectively. Finally, we demonstrate evolutionary conservation of hippocampal Sst-IN eGlyR expression and eGlyR- mediated SWR regulation in the nonhuman primate (NHP) hippocampus confirming functional availability of this target for therapeutic potential in higher mammalian species.

## Results

### eGlyR modulation of inhibitory drive and network synchronization in the early postnatal hippocampus

Detailed spatiotemporal profiling using multiplexed fluorescence in situ hybridization (mFISH) revealed remarkably high *Grin3a* expression within MGE-derived hippocampal interneurons, particularly CA1 Sst-INs, at early postnatal ages (∼postnatal day (P) 6-9) [Murillo 2021]. To initiate our study, we employed Nkx2.1^Cre/+^:Rpl22(RiboTag)^HA/HA^ (MGE-Ribotag) mice to independently validate and extend these findings. This Ribotag-based sequencing strategy provides a robust quantitative platform to examine mRNAs associated with the translational machinery for protein production selectively within the population of MGE-derived interneurons [Mahadevan 2020, Sanz 2009]. We observed an enrichment of *Grin3a* within hippocampal MGE-derived interneurons relative to bulk hippocampal tissue from early postnatal ages through adulthood (Figure 1A). Notably, bulk hippocampal *Grin3a* levels dramatically declined by 2 months of age, as expected given the well-established reduction in dorsal CA1 PNs [Ciabarra 1995, Sucher 1995, Sun 1998, Sasaki 2002, Zhang 2002, Perez-Otano 2016, Wee 2016, Murillo 2021, Crawley 2022, Bossi 2023]. However, MGE-derived interneurons retained high expression of *Grin3a* at all time points examined consistent with findings in adult neocortex [Pfeffer 2013, Paul 2017, Yao 2021, Bossi 2022]. Examination of dorsal hippocampal sections from a newly developed mClover- GluN3A reporter mouse line confirmed persistent GluN3A expression in a sparse population of cells outside the PN layer, prominently Sst+ neurons of stratum oriens, at adult stages when CA1 PN labeling is dramatically attenuated (Figure 1B).

**Figure 1.**
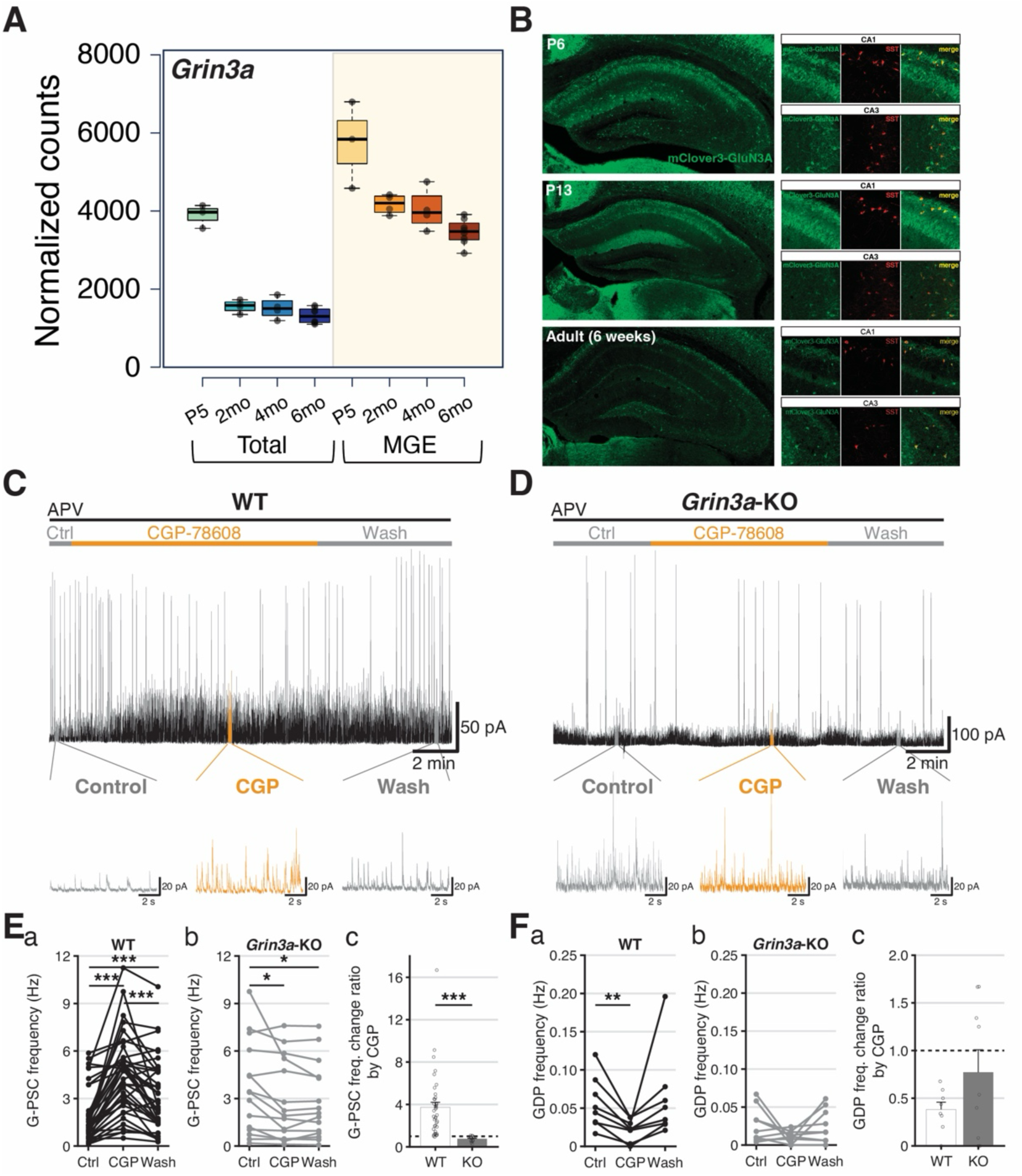
Developmental expression of *Grin3a* transcripts and G-PSC/GDP modulation by functional eGlyRs in early postnatal hippocampus. **A.** Developmental profile of *Grin3a* single RNA-seq data hippocampus using Ribo-tag. **B.** Developmental change of GluN3a expression images from mClover3-GluN1a mice demonstrating significant co-expression with somatostatin (Sst). **C-D.** Single trace example of spontaneous GABAergic postsynaptic currents (G-PSCs) and GDPs in CA3 pyramidal neurons (PNs) from WT (C), *Grin3a*-KO mouse brain slices (D). **E.** G-PSC frequency change in (a) WT, (b) *Grin3a*-KO, and (c) comparison of G-PSC frequency change ratio by CGP. **F.** Giant depolarizing potential (GDP) frequency change in WT (a), *Grin3a*-KO (b), and normalized comparison of GDP frequency change by CGP in WT and *Grin3a*-KO mice (c). (WT: 62 cells/slices from 26 mice, P4.5-8.5), *Grin3a*-KO: 32 cells/slices from 13 mice, P5.5-12.5 *pre- incubated in APV 100 μM.

At early postnatal ages the hippocampal network is dominated by GABAergic tone that exerts an excitatory influence upon postsynaptic PNs prior to establishment of low intracellular chloride levels by upregulation of KCC2 [Ben-Ari 2002, Ben-Ari 2007]. Between the ages of P5-10 this depolarizing GABAergic input coordinates spontaneous, synapse-driven network synchronizations in the form of giant depolarizing potentials (GDPs) that propagate throughout the hippocampus to drive activity-dependent circuit maturation and MGE-derived interneurons, particularly Sst-INs, are strongly implicated in this pacing [Ben-Ari 1989, Picardo 2011, Wester 2016, Flossmann 2019, Cossart 2022]. To probe for eGlyR-mediated regulation of GABAergic tone in the developing hippocampus we recorded spontaneous GABAergic postsynaptic currents (G-PSCs) and the GABAergic component of GDP-associated currents (GDP-Is) in acute hippocampal slices from P5.5–9.5 wild-type and *Grin3a* knockout (*Grin3a*-KO) mice while applying CGP-78608 to remove any eGlyR desensitization (Figure 1C-F). GABAergic synaptic events were electrically isolated by voltage-clamp at 0 mV where both G-PSCs and GDP-Is are detectable as outward GABA_A_R-mediated currents without significant contamination from phasic glutamatergic transmission (alpha-amino-3-hydroxy-5-methyl-4-isoxazolepropionic acid receptor (AMPAR)- mediated) needed in concert with GABAergic activity to sustain GDPs. Recordings were performed in CA3 PNs which lack *Grin3a* (Figure 1B; Supplemental Figure 1) to limit confounds of pyramidal cell eGlyRs through network effects and conventional NMDARs were blocked to eliminate confounds of CGP-78608 effects at their glycine binding sites. Under these conditions, CGP-78608 robustly increased CA3 PN G-PSC frequencies in WT but not *Grin3a*-KO mice in a reversible fashion (Figure 1C-E). Concomitantly, we consistently observed that GDP-I frequencies declined by the end of CGP-78608 application in WT mice without consistent effects in *Grin3a*- KOs (Figure 1F). Our initial expression profiling and functional observations suggest that immature hippocampal interneurons express eGlyRs that are liganded and desensitized by ambient Gly levels. Moreover, engaging interneuron eGlyRs has the capacity to dramatically increase their excitability to enhance GABAergic tone throughout the hippocampal circuit and potentially disrupt spontaneous rhythmicity.

### *Grin3a* expression in hippocampal NGFCs and Sst-INs yields functional eGlyRs throughout development that influence excitability

To obtain a more granular appreciation of *Grin3a* expression within specific interneuron subtypes we analyzed single-cell RNA sequencing (scRNA-seq) data from adult mice when interneuron transcriptional identities are well defined and segregated ([Harris 2018, Hodge 2019, Krienen 2020, Yao 2021, Mahadevan 2021]). Focusing on GABAergic interneurons from a multi-institute integrated scRNA-seq dataset (see Methods) we confirmed that most Sst-INs express high levels of *Grin3a* consistent with prior studies (Figure 2A;Supplemental Figure 1) [Pfeffer 2013, Paul 2017, Murillo 2021, Bossi 2022]. Indeed, *Grin3a* serves as a secondary molecular marker for Sst-INs and within these cells the *Grin3a* locus exhibits prominently open chromatin and low DNA methylation states that are indicative of actively transcribed genes promoting cell identity ([Pfeffer 2013, Yao 2021]). We also discovered previously unappreciated *Grin3a* expression across most NGFC subsets (Figure 2A; Supplemental Figure 1). Amongst NGFCs *Grin3a* is expressed in both Lamp5^+^/Lhx6^+^ MGE-derived subsets, also known as Ivy cells (IvyCs), and Lamp5^+^/Lhx6^-^ caudal ganglionic eminence (CGE)-derived subsets that are both well represented in the rodent hippocampus [Fuentealba 2008, Tricoire 2010, Tricoire 2011]. Consistent with these scRNA-seq profiles mClover-GluN3A labeled cells co-expressed Sst in stratum oriens and neuronal nitric oxide synthase (nNOS) in putative NGFCs that comprise the dominant interneuron subtype in stratum lacunosum moleculare (Figure 2B; Supplemental Figure 1).

**Figure 2.**
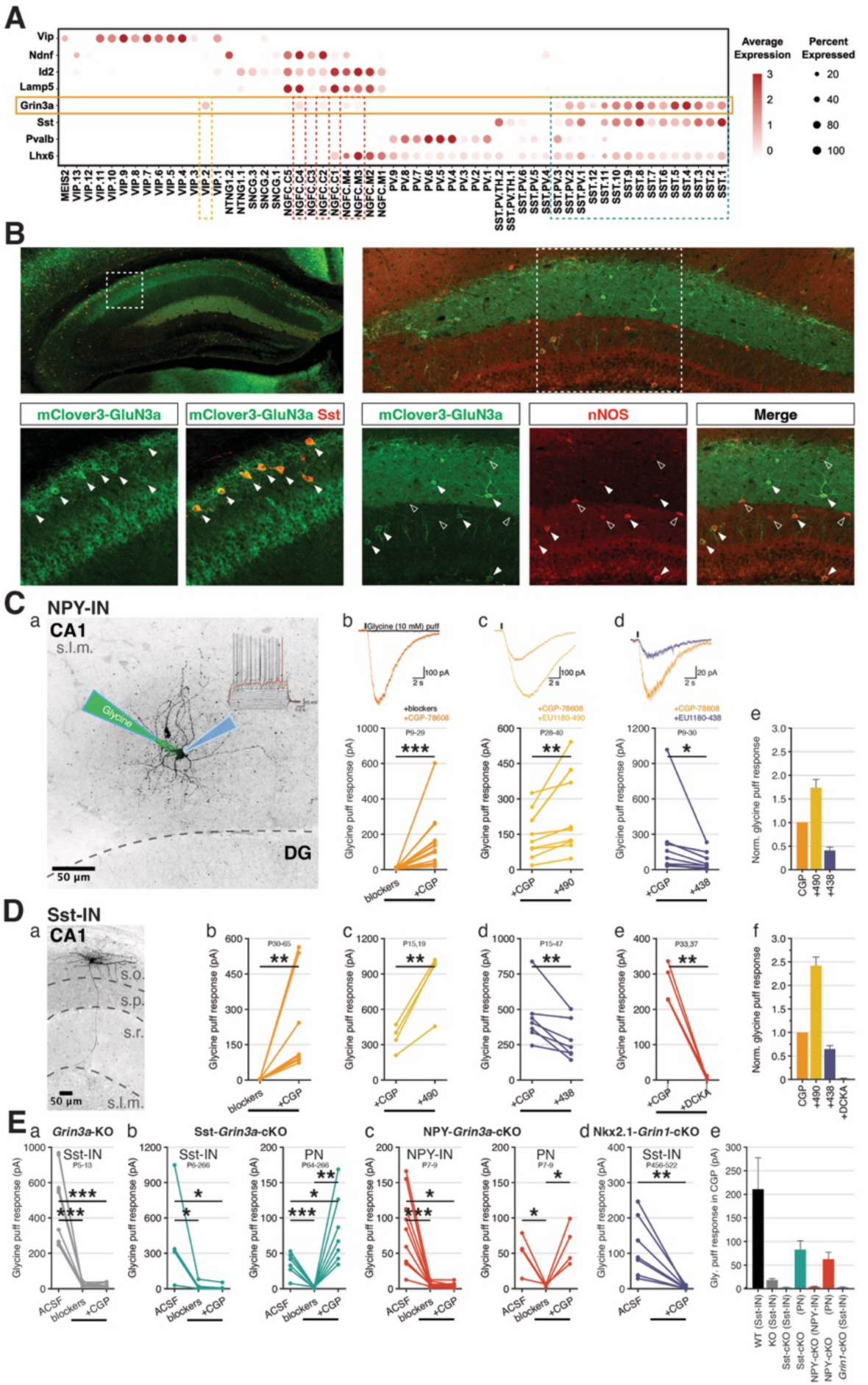
*Grin3a* expression and functional eGlyR-mediated currents in adult hippocampal INs. **A.** Survey of *Grin3a* expression across major GABAergic interneuron classes (Sst, somatostatin- expressing cells; PV, Parvalbumin-expressing cells; NGFC, neurogliaform cells; VIP, vaso-intestinal peptide-expressing cells) from mice cortex and hippocampus. **B.** Co-expression of *Grin3a* with Sst (left), and nNOS (right) respectively. **C.**(a) Whole cell patch-clamping an NPY-IN with glycine puff. Glycine puff response of NPY-IN in (b) desensitization reducer (CGP), (c) allosteric modulators positive (490), (d) negative (438), and (e) normalized glycine puff responses. **D.**(a) A sample image of Sst-IN (OLM-IN) (b) Glycine puff response of Sst-IN in CGP, (c-d) allosteric positive/negative modulators, respectively, (e) competitive eGlyR blocker (DCKA), and (f) normalized glycine puff responses. **E.** Glycine puff responses of (a) Sst-IN in *Grin3a*-KO mice, (b) Sst-IN and PN in Sst-*Grin3a*-cKO, (c) NPY-IN and PN in NPY-*Grin3a*-cKO, and (d) Sst-IN in Nkx2.1- *Grin1*-cKO, and (e) average glycine puff responses.

To directly confirm functional eGlyR expression within hippocampal interneurons we electrophysiologically measured their responses to brief, pressure applied Gly (10 mM) delivered locally via a glass micropipette (Figure 2C-D). Sst-INs and NGFCs were targeted using Sst-Cre:Ai14 and NPY-Cre:Ai14 tdTomato reporter mice respectively. To pharmacologically isolate eGlyR- mediated currents, we included a cocktail of receptor antagonists in the perfusing artificial cerebrospinal fluid (ACSF) to block conventional ionotropic glycine (strychnine, 1 μM) glutamate (DL-APV, 50 μM, DNQX, 10 μM), and GABA receptors (picrotoxin, 50 μM; bicuculline, 10 μM). With this cocktail any initial current induced by Gly in regular ACSF was essentially abolished indicating minimal eGlyR-mediated current availability (Figure 2C-D and see Supplemental Figure 2 for sequential antagonist example). However, application of CGP-78608 in the presence of the antagonist cocktail unmasked robust Gly-evoked currents in both Sst-INs and NGFCs consistent with mediation through eGlyRs that are desensitized under baseline conditions (Figure 2C-D). Importantly, these Gly-evoked currents required *Grin3a*, as they were absent in either constitutive or conditional *Grin3a* knockout mice (*Grin3a*-KO; Sst-*Grin3a*-cKO and NPY-*Grin3a*-cKO), though ventral CA1 PNs retained CGP-78608 sensitive Gly-evoked currents in both interneuron conditional KOs (Figure 2E). Similarly, recordings from stratum oriens Sst-INs in Nkx2.1-*Grin1*- cKOs lacked CGP-78608 sensitive Gly-evoked currents consistent with the requirement for both GluN1 and GluN3 to form functional eGlyRs (Figure 2E). Further pharmacological evidence that the currents observed in hippocampal interneurons reflect bona fide eGlyR conductance includes antagonism by the GluN1/3A Gly binding site antagonist 5,7-dichlorokynurenic acid (DCKA) and suppression by the negative allosteric modulator EU1180-438 (Figure 2C-D, [Chatterton 2002, Awobuluyi 2007, Zhu 2020, Bossi 2022]). In addition, we found that CGP-78608 unmasked Gly-evoked currents in both Sst-INs and NGFCs displayed significant potentiation in response to a newly developed GluN3-selective positive allosteric modulator of eGlyRs, EU1180-490, identified via focused library screening of human GluN1/GluN3A (Figure 2C-D, [Nguyen 2025]). In sum, these findings reveal robust eGlyR expression within hippocampal Sst-INs and NGFCs providing potential cellular mediators for the regulation of GABAergic tone onto CA3 PNs by CGP-78608 in the developing hippocampus. Indeed, a more detailed temporal developmental profiling of pressure applied Gly-evoked responses confirmed functional eGlyR expression within Sst-INs and NGFCs at early postnatal ages with Sst-INs exhibiting a modest reduction in current amplitudes with age and NGFCs displaying increased current amplitudes (Supplemental Figure 3).

Building on these findings, we next investigated whether ambient Gly working through eGlyRs has the capacity to directly modulate excitability of *Grin3a*-expressing hippocampal interneurons as suggested by the ability of CGP-78608 to drive GABAergic tone. First, we assessed whether eGlyRs provide a tonic current in hippocampal interneurons by examining the effects of DCKA on membrane holding/leak currents in the presence of the conventional ionotropic glycinergic, glutamatergic, and GABAergic receptor antagonist cocktail. Application of DCKA modestly reduced inward holding current at −70 mV in NGFCs but not Sst-INs (Figure 3A-B). This differential influence of tonic eGlyR currents in the two hippocampal interneuron subtypes is reminiscent of differences reported between amygdala principal cells and neocortical Sst-INs and may reflect regional and/or cell-type specific differences in ambient *ex vivo* Gly levels, as well as eGlyR redox and desensitization states [Grand 2018, Bossi 2022]. Although recorded in the same gross *ex vivo* structure under the same recording conditions NGFCs and Sst-INs were targeted in very deep or superficial CA1 respectively potentially with unique microenvironmental influences. Conversely, both NGFCs and Sst-INs consistently displayed an increase in holding current in response to application of CGP-78608 indicating sufficient endogenous ambient Gly levels to ligand and largely desensitize the eGlyRs on both interneuron populations (Figure 3C-D). The effects of CGP-78608 on holding current were reversed by DCKA and enhanced by EU1180-490 consistent with mediation through eGlyRs (Figure 3D and Supplemental Figure 4). Moreover, in the current-clamp recording configuration CGP-78608 depolarized both NGFCs and Sst-INs sufficiently to induce or increase action potential (AP) firing (Figure 3E-F). Interestingly, EU1180- 490 alone could also drive increased firing of Sst-INs, suggesting that this positive allosteric modulator may relieve eGlyR desensitization directly (Figure 3F). Together our findings conclusively demonstrate that ambient Gly levels have the capacity to strongly influence NGFC and Sst-IN excitability, and hence circuit GABAergic tone, throughout development via eGlyRs within these interneuron subpopulations.

**Figure 3.**
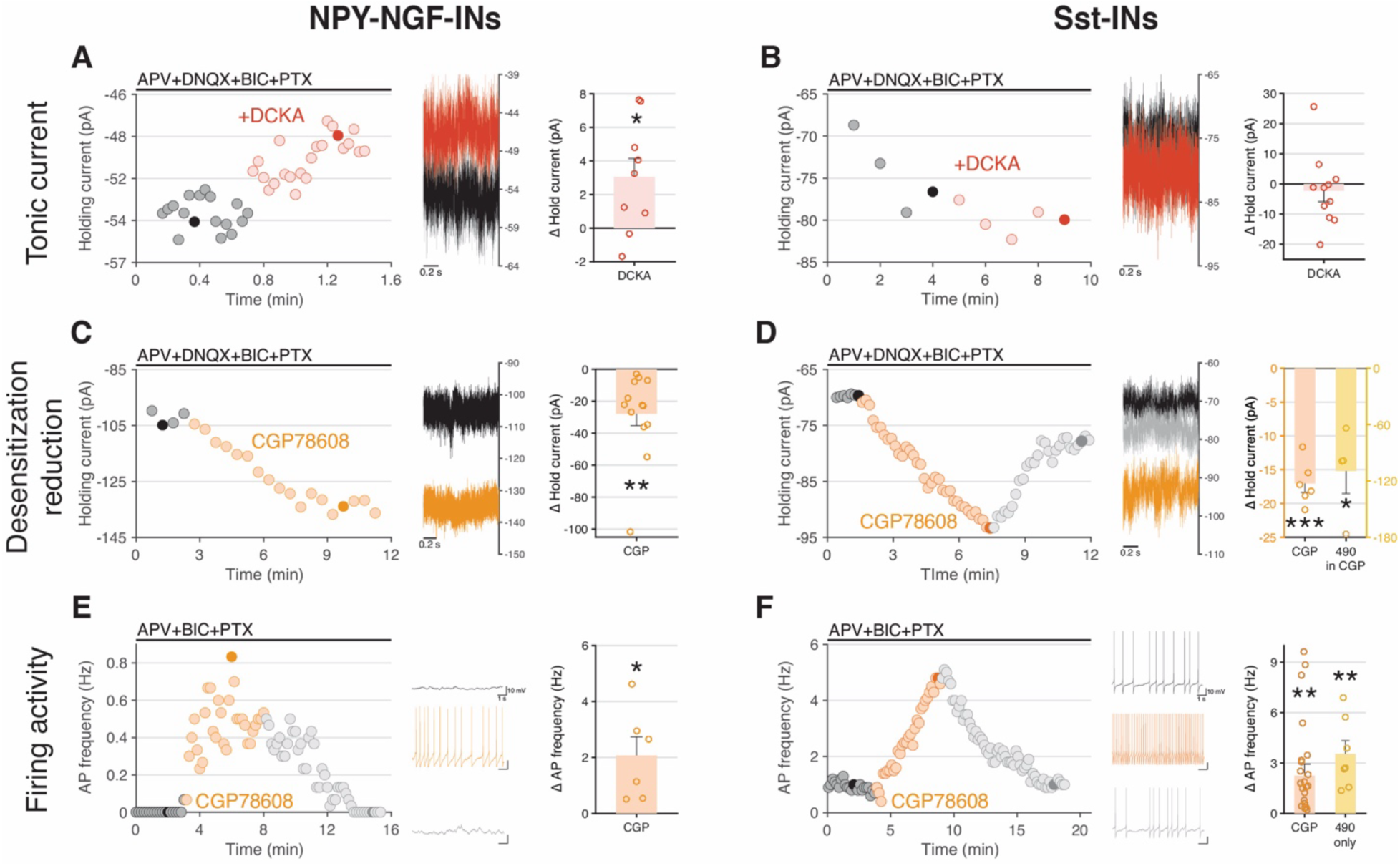
eGlyRs are tonically liganded by endogenous glycine in hippocampal SST and NPY- NGF-INs. **A-B.** eGlyR tonic current blocked by DCKA in NPY-INs, but no tonic current in Sst-INs. **C-D.** Holding current increase by desensitization reduction by CGP and/or EU1180-490 in NPY-INs and Sst-IN respectively. **E-F.** Firing activity increase by CGP or EU1180-490 in NPY-INs and Sst-INs respectively.

### An unexpectedly outsized contribution from NGFCs/IvyCs in early postnatal GABAergic tone and rhythmicity under control and CGP-78608 conditions

A significant body work has implicated Sst-INs as major contributors to early postnatal hippocampal and neocortical network GABAergic tone, coordination of population synchrony, and circuit connectivity/maturation [Picardo 2011, Anastasiades 2016, Marques-Smith 2016, Oh 2016, Tuncdemir 2016, Modol 2017, Flossmann 2019, Wang 2019, Su 2020, Cossart 2022, Dard 2022]. However, similar studies regarding the role of NGFCs/IvyCs in nascent network function are lacking despite these cells representing perhaps the most abundant dendrite targeting interneurons of the hippocampus [Bezaire 2013, Overstreet-Wadiche 2015]. To dissect the contributions of Sst-INs and NGFCs to GABAergic tone and pacing within the developing hippocampus under control and CGP-78608 conditions we employed an optogenetic strategy using conditional expression of the light-activated proton pump Achaerhodopsin-3 (Arch) to silence distinct interneuron populations (Figure 4). First, to determine the maximal effectiveness of this approach we used pan-GABAergic neuron Arch mice generated by driving conditional Arch expression with Cre under control of the vesicular inhibitory amino acid transporter gene (VGAT- Cre:ArCh). In control conditions (normal ACSF) light driven suppression of all GABAergic interneurons immediately and persistently blocked the majority of G-PSCs and GDP-Is recorded in early postnatal CA3 PNs of VGAT-Cre:ArCh mice (Figure 4A, C-F). Any residual G-PSCs and GDP- Is in light could reflect AP independent spontaneous GABA release and/or stochastic escapes from Arch-mediated suppression by interneurons with high input resistance during prolonged light epochs (for example, by a large spontaneous AMPAR-mediated glutamatergic input). The effects of Arch activation on G-PSC and GDP-I frequencies reversed upon light termination which typically yielded a bout of “rebound light-off GDPs” as expected for the synchronous release of interneurons from hyperpolarization that can drive them to AP threshold triggering a window of reverberant network activity due to the excitatory nature of GABA at this stage of development (Figure 4A, G and see [Wester 2016, Flossmann 2019]). Critically, light driven suppression of interneuron activity in VGAT-Cre:ArCh mice significantly inhibited G-PSCs in the presence of CGP- 78608 confirming that our optogenetic approach continues to restrain interneurons under conditions of eGlyR-mediated enhanced excitability (Figure 4B-D).

**Figure 4.**
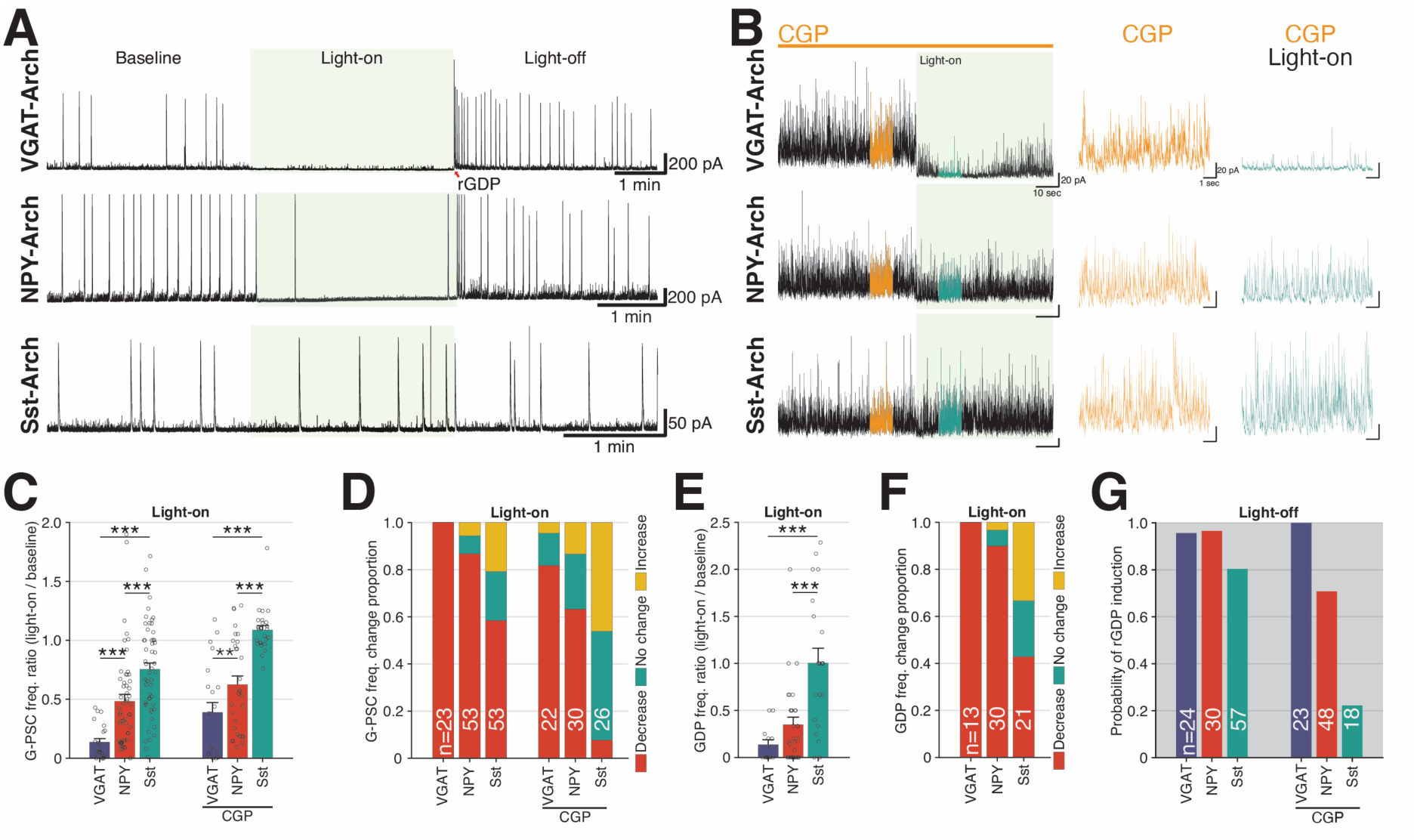
Relative contributions of NPY-NGF and SST INs in eliciting early hippocampal inhibition and network synchrony. **A.** Whole-cell recording samples of CA3-PN in brain slices from VGAT-ArCh (top), NPY-ArCh (middle), and Sst-ArCh (bottom) mice with light-stimulation. **B.** Examples of G-PSC change by light-on in presence of CGP-78608 in CA3-PN in hippocampal slices from VGAT-ArCh (top), NPY-ArCh (middle), and Sst-ArCh (bottom) mice **C.** G-PSC frequency change ratio by light-on in VGAT-Arch, NPY-Arch, and Sst-Arch mouse hippocampus. **D.** G-PSC frequency change proportion; decrease (red), change less than +- 10 % (green), and increase (yellow). **E.** GDP frequency change ratio by light-on. **F.** Light-on evoked GDP frequency change proportion; decrease (red), no change (green), increase (yellow). **G.** Probability of light-off- evoked rebound GDP (rGDP). (VGAT-Arch: 20 cells/slices from 5 mice, P6.5-8.5, NPY-Arch: 39 cells/slices from 10 mice, P5.5-8.5, Sst-Arch: 30 cells/slices from 10 mice, P5.5-8.5).

In control conditions, selective silencing of Sst-INs with light while recording from early postnatal CA3 PNs yielded highly inconsistent effects that on average modestly depressed G-PSC frequencies but did not significantly alter GDP-Is (Figure 4A, C-F). Successful light-mediated control of the Sst-IN population was evidenced by the high probability of light-off rebound GDPs observed in Sst-Cre:ArCh mice (Figure 4A, G). Surprisingly, Sst-IN silencing failed to significantly depress G-PSCs in the presence of CGP-78608, suggesting that the massive recruitment of GABAergic tone by eGlyRs occurs largely independent of Sst-IN output (Figure 4B-D). In contrast, optogenetic silencing in NPY-Cre:ArCh mice more closely matched our observations from pan- GABAergic neuron silencing with robust inhibition of G-PSC and GDP-I frequencies in control conditions, generation of rebound light-off GDP-Is, and continued inhibition of G-PSCs in the presence of CGP-78608 (Figure 4). Though NPY is strongly expressed in both NGFCs and Sst-INs, potentially confounding interpretation of our optogenetic results, subtractive reasoning points to an outsized role for NGFCs/IvyCs in providing GABAergic tone and rhythmicity within the early postnatal CA3 hippocampus. Indeed, the inability of selective Sst-IN silencing to match the impact of NPY-IN silencing on control G-PSCs and GDP-Is requires consideration of an NPY^+^/Sst^-^ IN cohort, with NGFCs/IvyCs being the most logical candidates. This interpretation is further bolstered by considering the limited possible sources of GABAergic input recruited by eGlyRs. As NPY-IN, but not Sst-IN, silencing significantly blocks G-PSCs recruited by eGlyR activation and *Grin3a* expression is largely limited to NGFCs/IvyCs and Sst-INs, our findings implicate NGFCs/IvyCs as the dominant source of CGP-78608 recruited GABAergic tone in the developing CA3 hippocampal circuit.

To independently test whether NGFCs/IvyCs are largely responsible for the increased GABAergic tone recruited by eGlyR activation in early postnatal tissue, we evaluated CGP-78608 effects on G-PSCs and GDP-Is in CA3 PNs of conditional *Grin3a* knockout mice (Figure 5). For these experiments we separated the CGP-78608 effects on G-PSCs and GDP-Is into early and late phases since closer temporal inspection of our recordings in wild type mice revealed that while G-PSC frequencies continually increase and plateau during drug application, GDP-Is undergo biphasic effects first increasing in frequency then decreasing (Figure 5). The initial increase in GDP-I frequency is accompanied by decreasing amplitudes until ultimately events cannot be reliably resolved from the increased G-PSCs (eg. Figure 5A-B). This profile for GDP-Is is consistent with the ability of CGP-78608 to increase IN excitability and ultimately drive them beyond threshold (Figure 3). A transient increase in GDP frequency could reflect enhanced excitability of INs participating in GDP generation and propagation; however, as greater numbers of INs are driven to threshold out of phase with the rhythmic network, and perhaps even into depolarization block, the ability to sustain synchronization may be compromised prompting degradation of GDP-Is into uncoordinated unitary events. Importantly, in constitutive *Grin3a*-KO mice all early and late phase CGP-78608 effects on G-PSCs and GDP-Is are eliminated confirming the pharmacological specificity for *Grin3a*-containing receptors in mediating transient and sustained changes in GABAergic tone and circuit pacing (Figure 5Ca,b,f-Da,b,f). However, selective removal of *Grin3a* across the entire IN population (VGAT-*Grin3a*-cKO mice) eliminated only the CGP-78608 induced increase in G-PSCs and the transient increase in GDP-Is (Figure 5Cc,f-Dc,f). Thus, late-stage GDP inhibition by CGP-78608 could reflect network changes triggered by eGlyRs in PNs, for example through polysynaptic interactions between eGlyR expressing CA1 PNs to the CA3 network that enhance network excitability to disrupt rhythmicity [Sik 1994, Kim 2015, Bocchio 2020]. Surprisingly, but consistent with our optogenetic findings, selective removal of *Grin3a* from Sst- INs had no effect on early or late phase effects of GCP-78608 on CA3 PN G-PSCs or GDP-Is (Figure 5Ce,f-De,f). In contrast, NPY-*Grin3a*-cKO mice failed to exhibit persistent or transient increases in CA3 PN G-PSCs or GDP-Is respectively in response to CGP-78608, essentially phenocopying the VGAT-*Grin3a*-cKO mice (Figure 5Cd,f-Dd,f). These findings are entirely consistent with our optogenetic results further indicating that the majority of GABAergic recruitment onto CA3 PNs by removal of eGlyR desensitization arises from NGFCs/IvyCs highlighting a previously unappreciated critical role for this class of interneuron in nascent hippocampal circuit activity.

**Figure 5.**
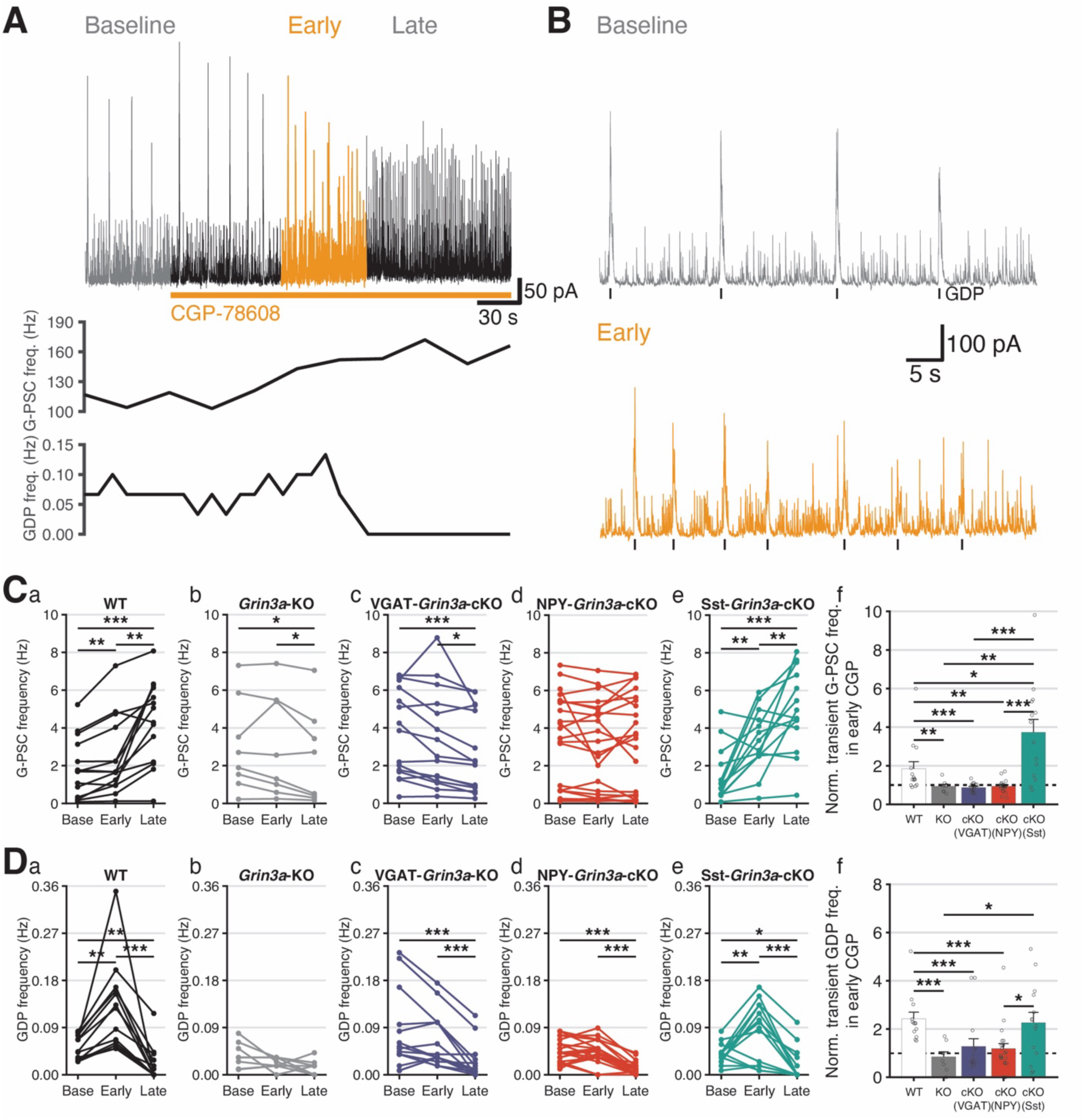
Recruitment of GABAergic tone and GDP generation is predominantly mediated by activation of eGlyRs expressed on NPY-NGF-INs. **A.** An example recording of GDP with CGP- 78608 (upper), G-PSC frequency (middle), and GDP frequency (bottom). **B.** Enlarged traces in baseline (upper)/CGP-Early(bottom) phases from A. **C.** G-PSC frequency change before, immediate (early) and stable (late) phase of CGP application in (a) WT, (b) *Grin3a*-KO, (c) VGAT- *Grin3a*-cKO, (d) NPY-*Grin3a*-cKO, (e) Sst-*Grin3a*-cKO mouse hippocampal slices, and (f) normalized immediate G-PSC frequency change by CGP. **D.** GDP frequency change before, early and late CGP application in (a) WT, (b) *Grin3a*-KO, (c) VGAT-*Grin3a*-cKO, (d) NPY-*Grin3a*-cKO, (e) Sst-*Grin3a*-cKO mouse hippocampal slices, and (f) normalized immediate GDP frequency change by CGP.

### eGlyR-mediated excitation of Sst-INs regulates sharp wave ripples in adult hippocampal circuits

In the mature hippocampus the dominant spontaneous synchronous network signature is that of sharp wave ripples (SWRs) that are typically observed *in vivo* during “off-line” brain states such as quiet wakefulness, consummatory behaviors, and non-REM sleep [Buzsáki 2015]. These high-frequency synchronized oscillatory events play a critical role in memory consolidation and cognition by allowing rapid network replay of exploratory driven activity patterns over compressed time periods and are characterized by large-amplitude sharp waves with superimposed high-frequency ripple oscillations [Girardeau 2009, Jadhav 2012, Buzsáki 2015, Joo 2018]. SWRs are temporally and spatially coordinated by stereotyped activation and silencing of discrete interneuron subtypes, including Sst-INs with oriens-lacunosum moleculare (OLM) Sst- INs exhibiting suppression or excitation depending on experimental preparation and long-range projecting Sst-INs undergoing strong recruitment during SWRs [Royer 2012, Varga 2012, Somogyi 2014, Buzsáki 2015, Katona 2017, Szabo 2022]. Interestingly, interference with *Grin3a* expression can disrupt cognition and consolidation of episodic memories; however, the influence of eGlyRs on mature hippocampal network oscillations has not been previously investigated [Gallinat 2007, Roberts 2009, Papenberg 2014]. As our findings confirm continued expression of functional eGlyRs in adult hippocampal Sst-INs we directly evaluated whether eGlyRs have the capacity to influence SWR rhythmicity (Figure 6).

**Figure 6.**
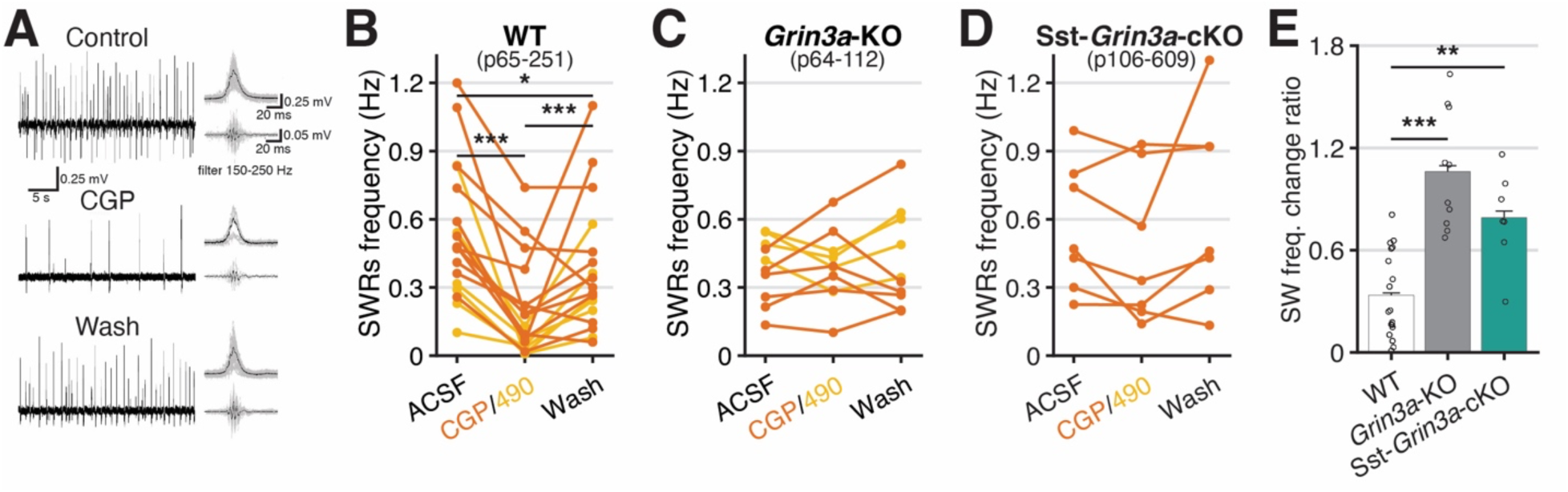
Activation of eGlyRs shape adult hippocampal sharp wave ripple dynamics. **A.** An example of sharp wave ripples. **B-D.** Sharp wave frequency change by CGP-78608/EU1180-490 in WT, *Grin3a*-KO, and Sst-*Grin3a*-cKO, respectively. **E.** SPW change rate in drug normalized by frequency in ACSF. All recordings were performed with APV (50 μM) in bath solution.

Spontaneous SWRs were readily detected with local field potential (LFP) recordings in the CA3 PN layer of acute hippocampal slices from adult mice maintained at interface (Figure 6A, [Hajos 2009, Pelkey 2015]). Importantly, spontaneous SWRs persist, and may even be enhanced, in the absence of conventional NMDAR-mediated transmission, and thus, our recordings were performed with conventional NMDARs blocked to eliminate pharmacological confounds to data interpretation of findings with eGlyR-related compounds [Behrens 2005, Colgin 2005]. In slices from wild type mice application of CGP-78608 or EU1180-490 significantly depressed the frequency of SWR occurrence in a reversible fashion (Figure 1A-B,E). Inhibition of SWRs by CGP- 78608 or EU1180-490 was absent in *Grin3a*-KO mice confirming the pharmacological specificity of these reagents for eGlyRs (Figure 6C,E). Most interestingly, selective removal of *Grin3a* from Sst-INs in Sst-*Grin3*-cKO mice greatly attenuated SWR sensitivity to CGP-78608 compared to control animals leaving modest inhibition that was not significantly different from data obtained in *Grin3a*-KOs (Figure 6D-E). Collectively, these findings illustrate that eGlyRs, particularly those expressed by Sst-INs, have the capacity to influence coordinated network rhythmicity in the adult hippocampus.

### Evolutionary conservation of eGlyR function in hippocampal interneurons

Genetic mutations or altered expression levels of *GRIN3A* in humans are implicated in cognitive deficits and diverse neurological/neuropsychiatric conditions such as schizophrenia, bipolar disorder, addiction, epilepsy, and Huntington’s Disease [Mueller 2004, Gallinat 2007, Marco 2013, Yuan 2013, Yang 2015, Lee 2016, Huang 2017, Marco 2018, Crawley 2022]. Historically, interpretation and posited roles of dysregulated *GRIN3A* in disease progression have focused on potential influences over conventional NMDARs (eg. triheteromeric GluN1/2/3 contributions) and phasic synaptic transmission [Kehoe 2013, Perez-Otano 2016, Crawley 2022]. The emerging picture in mice that GluN3A readily assembles as eGlyRs in diverse neurons expressing *Grin3a* to tonically regulate excitability through ambient Gly provides a new lens through which to consider GluN3A-associated disease etiology [Bossi 2023]. However, this demands functional confirmation that eGlyRs are evolutionarily conserved regulators of neuronal excitability. Thus, in our final series of experiments we aimed to validate our rodent findings in higher species by functionally interrogating hippocampal interneurons and network rhythmogenesis in adult non-human primates (NHPs).

Analysis of publicly available scRNA-seq data sets online indicate conserved *GRIN3A* expression within neocortical/hippocampal Sst-INs and NGFCs in both NHPs and humans (Supplemental Figure 5). By mFISH we confirmed that *GRIN3A* exhibits a high degree of colocalization with *SST* and *LAMP5* throughout the adult NHP (rhesus macaque) hippocampus (Figure 7A-B). To facilitate Sst-IN identification for functional evaluation of eGlyRs we intracranially injected adult macaques with novel enhancer based viral tools designed to limit transgene expression to Sst-INs, pAAV-PHP.eB-BiSSTe10-dTomato and pAAV-PHP.eB-BiSSTe4 [Furlanis 2025]. Then 5-6 weeks after hippocampal AAV introduction we made acute hippocampal slices and targeted fluorescently labeled cells for electrophysiological interrogation (Figure 7C). Consistent with our mouse findings NHP Sst-INs exhibited CGP-78608 sensitive Gly-evoked excitatory currents in the cocktail of conventional glutamate, GABA, and glycine receptor antagonists (Figure 7D). Importantly, these Gly-evoked currents were facilitated by EU1180-490 consistent with mediation by eGlyRs (Figure 7D). Parallel recordings in NHP anterior hippocampal CA1 PNs similarly yielded CGP-78608 and EU1180-490 sensitive Gly-evoked currents (Figure 7D). Finally, as SWRs are evolutionarily conserved network oscillations observed across multiple species, including NHPs and humans [Buzsáki 2015] we assessed whether engaging eGlyRs by removal of desensitization/allosteric enhancement could regulate rhythmicity in NHPs. Congruent with our rodent findings, we found that either CGP-78608 or EU1180-490 significantly depressed the incidence of SWRs recorded in the CA3 PN layer of adult NHP slices in a reversible fashion (Figure 7E). In total our expression and functional profiling confirm strong conservation of eGlyR function over approximately 70-100 million years of evolution validating its relevance as a potential novel therapeutic target to regulate neuronal excitability across a constellation of human neurological and neuropsychiatric disorders.

**Figure 7.**
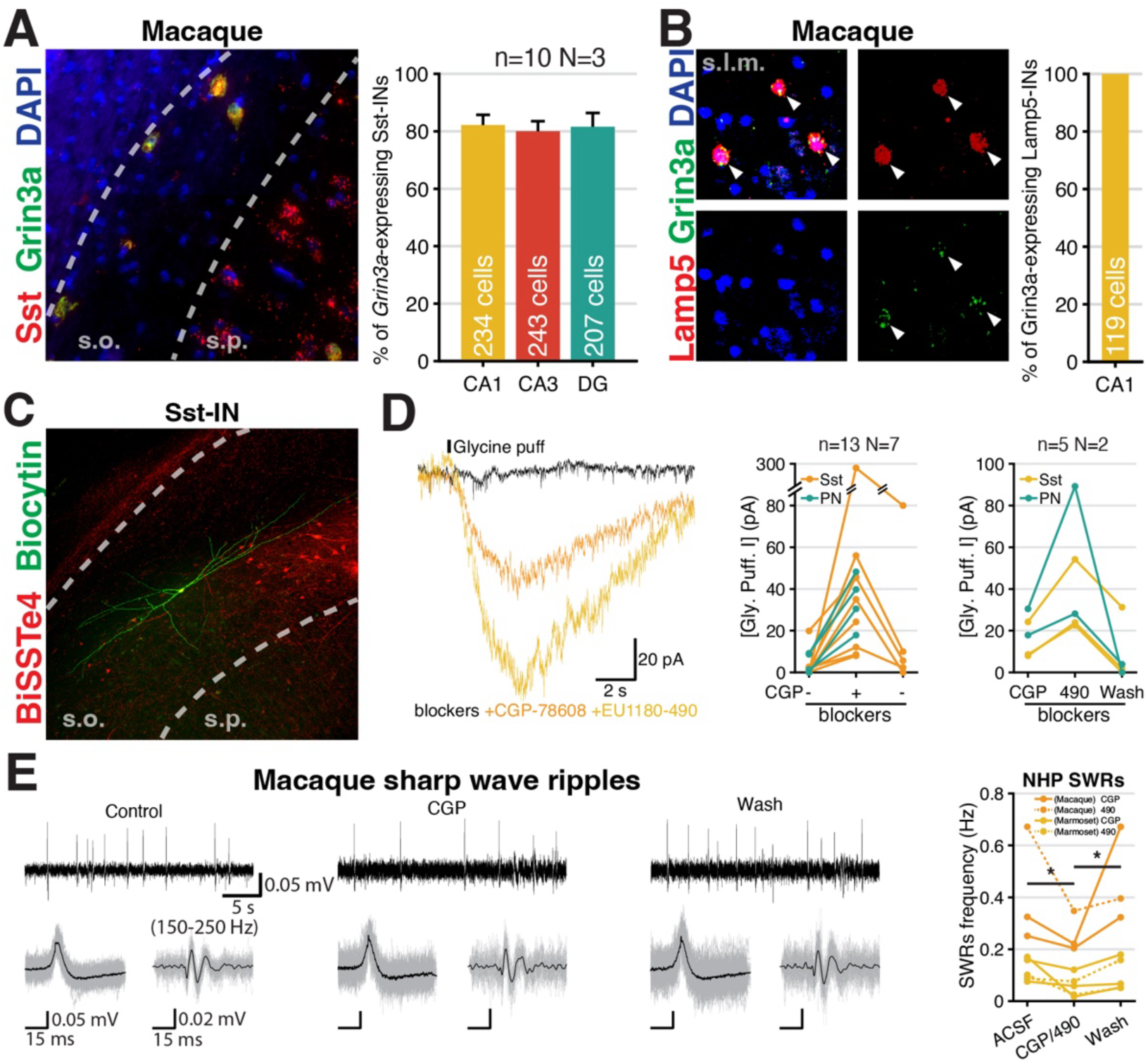
Evolutionary conservation of hippocampal interneuron eGlyR functional expression and role in network rhythmogenesis. **A.** Co-expression of Sst and *Grin3a* in a Macaque CA3 (stratum oriens) slice. **B.** Co-expression of Lamp5 and *Grin3a*. **C.** An example image of Sst-IN reported by pAAV-PHP.eB-BiSSTe4-dTomato (red) and post-hoc cell morphology recovery (green) in Macaque CA1 hippocampus. **D.** Glycine puff response of Sst-INs and PNs with CGP-78608 (middle) and CGP-78608 + EU1180-490 (right) in macaque hippocampus. **E.** SWRs frequency change by CGP-78608 / EU1180-490 in macaque (n=3, N=2) and marmoset (n=5, N=2). (n: number of slices, N: number of animals)

## Discussion

Emerging evidence indicates that GluN3A subunits frequently assemble with GluN1 to form unconventional NMDARs that serve as eGlyRs in diverse neurons throughout the brain, consistent with initial cloning studies [Perez-Otano 2001, Chatterton 2002, Grand 2018, Otsu 2019, Zhu 2020, Bossi 2022, Rouzbeh 2023, Pizzamiglio 2025]. Our present findings unequivocally demonstrate that *Grin3a*^+^ hippocampal Sst-INs and NGFCs/IvyCs express functional eGlyRs from early postnatal to adult ages and across evolution from mouse to macaque. Excitation of NGFCs/IvyCs and Sst-INs with endogenous ambient glycine through eGlyRs has the capacity to regulate spontaneous synchronized hippocampal network activities throughout development. Putative roles for GluN3A in synapse development and circuit function inferred from preclinical knockout/overexpression studies and genetic mutation linkage clinical studies are often modeled on cell autonomous influences of GluN3A over conventional NMDARs through formation of triheteromeric receptors or by serving as dominant negative subunits. Our findings, combined with others, highlight that eGlyR mediated regulation of cell and circuit excitability through both cell autonomous and cell non-autonomous mechanisms must also be considered to understand the role of *Grin3a*/GluN3A in brain development, plasticity, and disease [Otsu 2019, Bossi 2022, Hurley 2024, Pizzamiglio 2025].

The gating of eGlyRs is unusual with glycine serving as both agonist through high affinity GluN3A binding and functional antagonist by driving profound desensitization upon binding the lower affinity GluN1 subunit [Chatterton 2002, Yao 2006, Awobuluyi 2007, Madry 2007, Bossi 2022]. Thus, eGlyRs are well suited to provide tonic excitation at low levels of ambient glycine with an auto-inactivation mechanism at higher glycine levels to limit receptor mediated hyperexcitability/excitotoxicity. Interestingly, higher ambient glycine levels leading to GluN1 binding will promote a switch to increased phasic glutamatergic excitation by saturating co- agonist GluN1 binding sites of conventional NMDARs concomitant with the suppression of tonic eGlyR mediated excitation. Though we observed a modest influence of tonic eGlyR activation on NGFC resting membrane potential, the robust depolarizing influence of CGP-78608 on both Sst- INs and NGFCs indicates that ambient glycine levels throughout our acute hippocampal slices are by and large sufficient to bind GluN1 subunits and drive eGlyR desensitization. This is consistent with decades of electrophysiological interrogation of conventional NMDARs in acute hippocampal slices across many labs without the need for extracellular glycine supplementation. Our findings mirror observations in amygdala and neocortical Sst-INs that exhibit eGlyR desensitization under basal slice conditions [Bossi 2022]. In contrast, under the same conditions eGlyRs in amygdala PNs exhibit minimal desensitization and provide significant tonic depolarizing influence that is titrated through neuromodulation of glycine transporter activity [Bossi 2022]. Thus, even within the same gross brain structure under the same conditions eGlyR glycine-binding site occupancies may differ across neuronal subtypes according to differential extracellular ambient glycine levels. A very recent report provided evidence for tonic Gly liganding and depolarizing influence of eGlyRs on interneurons with somatic localization to stratum oriens but not PNs and interneurons residing in CA1 stratum pyramidale [Pizzamiglio 2025]. While these findings do not entirely align with our observations, perhaps due to recording conditions and animal ages, they further highlight potential for local Gly microenvironments in modulating cell excitability through eGlyRs. It will be important in future investigations to probe both sources and spatiotemporal plasticity of ambient hippocampal glycine *in vivo* across development. In addition, the physiological conditions and extent to which Sst-IN and NGFC/IvyC eGlyR functions are subject to activity dependent modulation through redox regulation, acidification, and zinc should also be evaluated [Madry 2007, Cummings 2017]. Moreover, it remains possible that there exists an as yet undiscovered endogenous neuroactive molecule that promotes eGlyR function through removal of desensitization or allosteric potentiation mechanistically analogous to CGP-78608 and 1180- 490.

GABAergic INs throughout the neocortex, amygdala, and hippocampus have common developmental origins in the ganglionic eminences and discrete IN subtypes in each of these structures display considerable genetic, physiological, and synaptic connectivity/function homologies [Zeisel 2015, Tremblay 2016, Pelkey 2017, Gouwens 2020, Hajos 2021, Yao 2021]. However, granular investigation can reveal regional differences in receptor expression profiles for homologous IN subpopulations across brain regions [Caccavano 2025]. Expression of eGlyRs within Sst-INs is universal across the cortical structures thus far examined ([Bossi 2022, Pizzamiglio 2025], and present findings). In contrast, parvalbumin (PV) IN expression of eGlyRs appears inconsistent with functional expression demonstrated in amygdala and hippocampal PV INs but not their neocortical counterparts [Bossi 2022, Pizzamiglio 2025]. Curiously, *Grin3a* is minimally expressed within transcriptionally defined rodent hippocampal and neocortical PV INs, though increases through evolution (Supplemental figure 5). These findings highlight the importance of careful cell-type specific evaluation across brain regions and species in mapping eGlyR expressing neurons to understand their circuit influences and potential for therapeutic leverage. In addition to the utility of emerging nervous system wide transcriptional data sets across species, identification of rodent GluN3A/eGlyR expressing neurons throughout the brain will be greatly facilitated by the newly developed mClover-GluN3A reporter mouse line.

Here we further identify hippocampal MGE- and CGE-derived NGFC/IvyC cohorts as neuronal elements through which ambient Gly can regulate circuit function by eGlyR signaling. Transcriptional profiling indicates conserved strong *Grin3a* expression levels in neocortical NGFCs defined by *Id2* and *Lamp5* expression (Supplemental Figure 5A). In evaluating NGFCs/IvyCs through the lens of eGlyRs we discovered a previously unappreciated capacity for these interneurons to dramatically shape GABAergic tone and network pacing in the developing hippocampus. Past studies have largely implicated Sst-INs as major contributors to early postnatal hippocampal and neocortical network GABAergic tone, coordination of population synchrony, and circuit connectivity/maturation [Picardo 2011, 1, Marques-Smith 2016, Oh 2016, Tuncdemir 2016, Modol 2017, Flossmann 2019, Wang 2019, Su 2020, Cossart 2022, Dard 2022]. However, while synchronization of Sst-IN activation has the capacity to generate GDP-like events (eg. light off events in Sst-Cre: ArCh) minimizing Sst-IN activity fails to consistently depress spontaneous GABAergic tone or intrinsic GDP pacing in the developing hippocampus (see also [Flossmann 2019]). In contrast, the same optogenetic strategy to silence NGFC/IvyC activity consistently suppressed spontaneous GABAergic tone and GDPs in CA3 PNs. Moreover, NGFC/IvyC inputs were found to dominate the excess inhibitory drive recruited by eGlyR activation with CGP-78608 in the immature hippocampus. These findings, together with recent evidence implicating immature parvalbumin interneurons in early network GABAergic pacing [Caccavano 2025], highlight the need for further investigation of multiple cardinal IN classes in coordinating developing hippocampal circuit dynamics.

The increasing availability of publicly accessible single cell transcriptomic data sets has greatly facilitated genetic profiling of discrete neuronal populations across species as a starting point for validating translational relevance of rodent centric studies to higher species. However, opportunities for functional validation in neuronal cells and circuits of higher species remains limited by tissue availability and genetic identification of discrete cell types. Here we employed recently developed enhancer based AAV tools to facilitate selective targeting of Sst-INs in live NHP tissue for direct functional interrogation of the evolutionary conservation of eGlyR expression in this neuronal subpopulation [Furlanis 2025]. Further development of such tools for cell-type specific targeting will greatly expand the discrete neuronal subtypes throughout the brain for targeting across species in preclinical studies requiring selective cell labeling/actuation and for future therapeutic intervention or gene therapy [Tasic 2025]. Our findings confirmed that expression of *GRIN3A* is indeed predictive of operational eGlyRs in NHPs as in rodents. The emerging picture from our current findings combined with others [Grand 2018, Zhu 2020, Bossi 2022], necessitate consideration of eGlyR dysregulation in addition to conventional NMDARs as an underlying feature of neurological disorders associated with genetic mutations or alterations in *GRIN3A* expression such as schizophrenia, bipolar disorder, addiction, epilepsy, and Huntington’s Disease [Mueller 2004, Gallinat 2007, Marco 2013, Yuan 2013, Yang 2015, Lee 2016, Huang 2017, Marco 2018, Crawley 2022]. Importantly, NHP hippocampal Sst-IN and CA1 PN eGlyRs displayed similar pharmacological profiles to those in rodents with CGP-78608 “awakening” the receptors from desensitization and the newly described allosteric modulator EU1180-490 able to potentiate their responses. Moreover, as in rodents, both compounds worked independent of conventional NMDARs to depress *in vitro* SWR activity, illustrating the ability to leverage eGlyRs expressed in circuit minority cell populations for regulation of network population activities across species. These findings provide a foundation for further native eGlyR evaluation across species in health and disease and confirm their potential as promising therapeutic targets using small molecule selective allosteric modulators.

## Acknowledgements

This research was supported [in part] by the Intramural Research Program of the National Institutes of Health (NIH). The contributions of the NIH author(s) are considered Works of the United States Government. The findings and conclusions presented in this paper are those of the author(s) and do not necessarily reflect the views of the NIH or the U.S. Department of Health and Human Services. We thank Apratim Mitra, Mira Sohn and Ryan Dale (NICHD Bioinformatics and Scientific Programming Core) for assistance with downloading and analyzing scRNAseq datasets. These analyses utilized the computational resources of the NIH Biowulf cluster (http://hpc.nih.gov).

## Author contributions

JHK was responsible for project conceptualization, experimental design, electrophysiology experiments, and imaging, analyses. KAP and CJM conceptualized, designed, and supervised the project. JHK, AV GAV, TB, RC, and KAP contributed to electrophysiology experiments. AV and GAV conducted AAV injections and surgery for mice. XY and SH conducted IHC, ISH, morphological reconstructions, and imaging. VM and APV conducted RNAseq analyses, RiboTag experiments and analysis. OCC, AIN and IPO provided Sst-Cre::*Grin3a*^fl/fl^ and mClover-GluN3A reporter IHC analysis. JHK, DA, KP generated Npy-/VGat-Cre:: *Grin3a*^fl/fl^ mice and maintained mouse colonies. DL, ZZ, HY, SFT, SLS, MPE, and DCL provided GluN3a specific allosteric modulators (EU1180-490 and EU1180- 438). MAGE, ACC, ACT, PH, AP, AM, JP, and BBA conducted NHP handling, injections, and surgeries. EF, YW, MD, BLG, JD, and GF provided support and design of enhancer AAV constructs. All authors participated in drafting the manuscript with JHK and KAP serving as principal writers. MAGE, BBA, GF, IPO, SFT and CJM provided critical funding support for the study.

## STAR★METHODS

### KEY RESOURCES TABLE

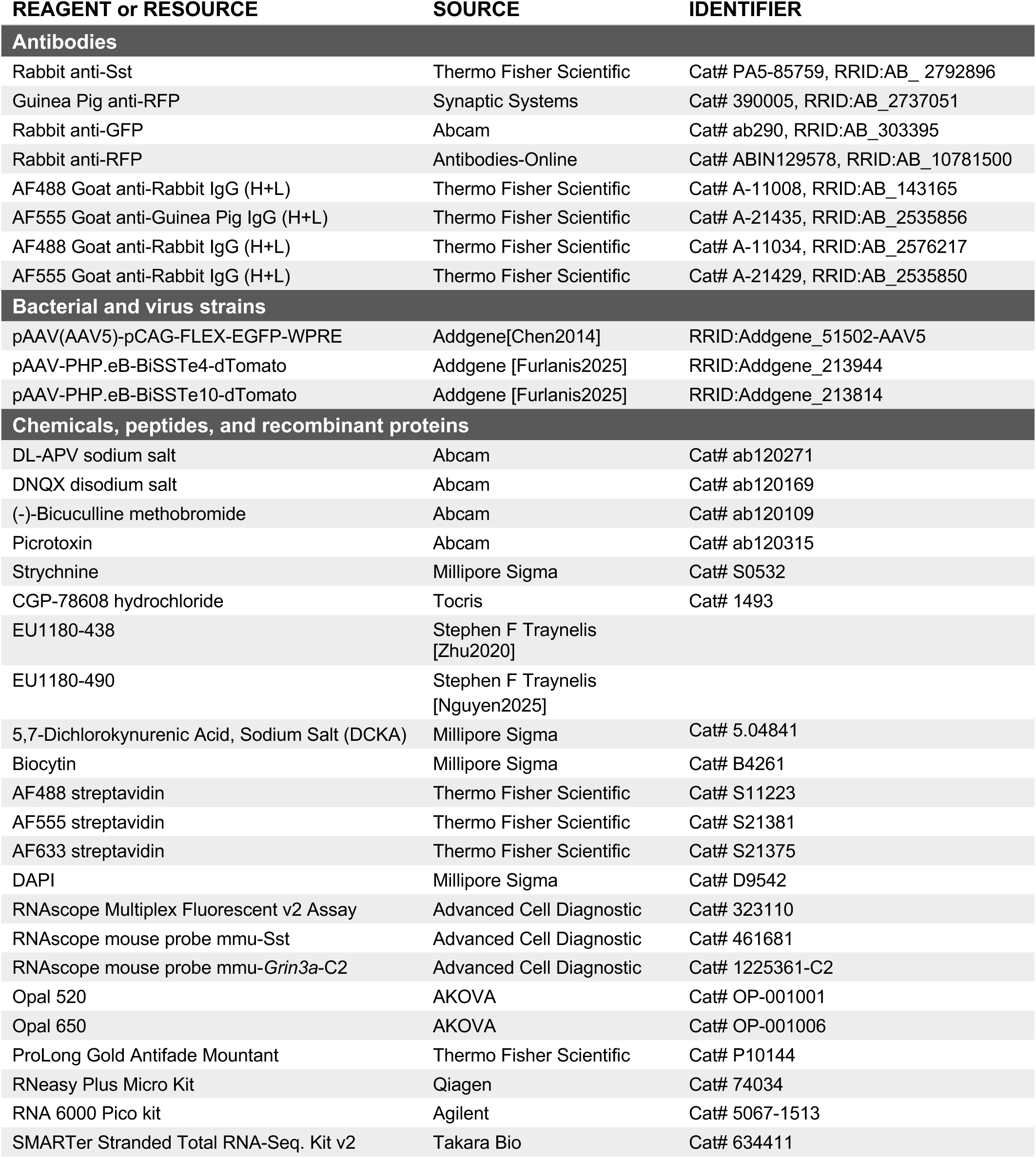
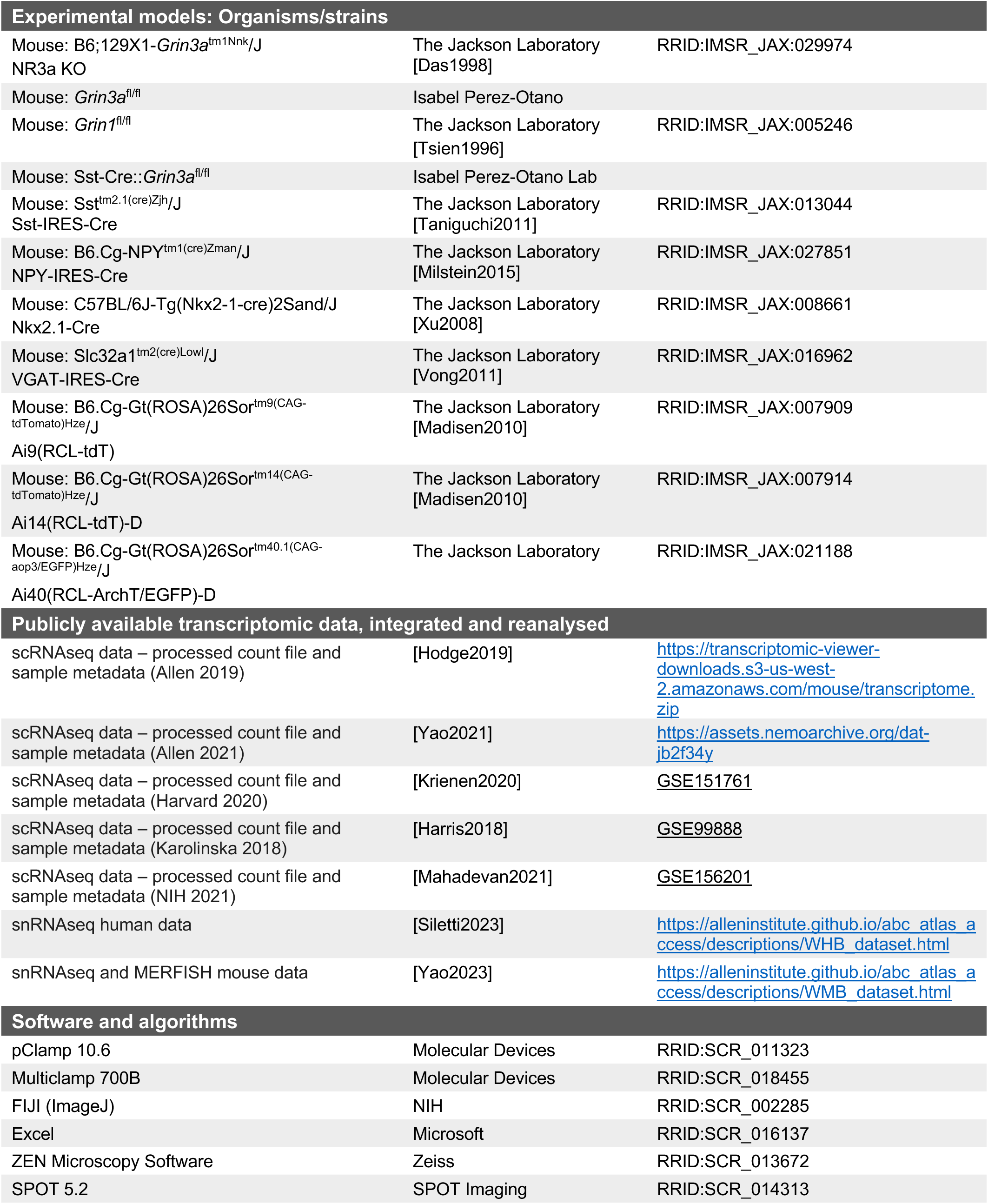
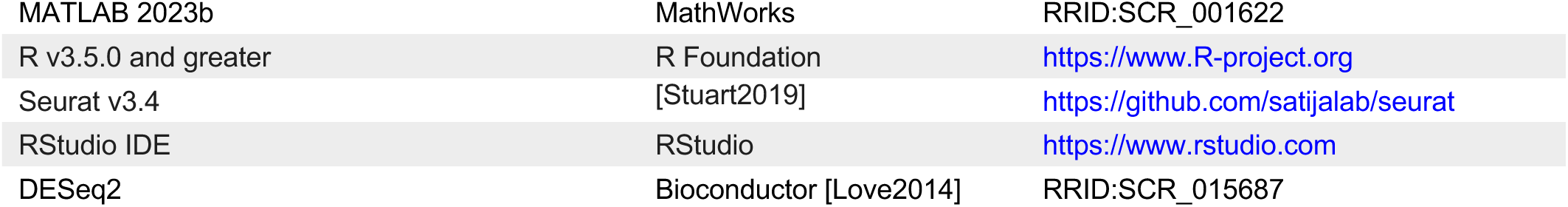

| REAGENT or RESOURCE | SOURCE | IDENTIFIER |
| --- | --- | --- |
| **Antibodies** |  |  |
| Rabbit anti-Sst | Thermo Fisher Scientific | Cat# PA5-85759, RRID:AB_2792896 |
| Guinea Pig anti-RFP | Synaptic Systems | Cat# 390005, RRID:AB_2737051 |
| Rabbit anti-GFP | Abcam | Cat# ab290, RRID:AB_303395 |
| Rabbit anti-RFP | Antibodies-Online | Cat# ABIN129578, RRID:AB_10781500 |
| AF488 Goat anti-Rabbit IgG (H+L) | Thermo Fisher Scientific | Cat# A-11008, RRID:AB_143165 |
| AF555 Goat anti-Guinea Pig IgG (H+L) | Thermo Fisher Scientific | Cat# A-21435, RRID:AB_2535856 |
| AF488 Goat anti-Rabbit IgG (H+L) | Thermo Fisher Scientific | Cat# A-11034, RRID:AB_2576217 |
| AF555 Goat anti-Rabbit IgG (H+L) | Thermo Fisher Scientific | Cat# A-21429, RRID:AB_2535850 |
| **Bacterial and virus strains** |  |  |
| pAAV(AAV5)-pCAG-FLEX-EGFP-WPRE | Addgene[Chen2014] | RRID:Addgene_51502-AAV5 |
| pAAV-PHP.eB-BiSSTe4-dTomato | Addgene [Furlanis2025] | RRID:Addgene_213944 |
| pAAV-PHP.eB-BiSSTe10-dTomato | Addgene [Furlanis2025] | RRID:Addgene_213814 |
| **Chemicals, peptides, and recombinant proteins** |  |  |
| DL-APV sodium salt | Abcam | Cat# ab120271 |
| DNQX disodium salt | Abcam | Cat# ab120169 |
| (-)-Bicuculline methobromide | Abcam | Cat# ab120109 |
| Picrotoxin | Abcam | Cat# ab120315 |
| Strychnine | Millipore Sigma | Cat# S0532 |
| CGP-78608 hydrochloride | Tocris | Cat# 1493 |
| EU1180-438 | Stephen F Traynelis [Zhu2020] |  |
| EU1180-490 | Stephen F Traynelis [Nguyen2025] |  |
| 5,7-Dichlorokynurenic Acid, Sodium Salt (DCKA) | Millipore Sigma | Cat# 5.04841 |
| Biocytin | Millipore Sigma | Cat# B4261 |
| AF488 streptavidin | Thermo Fisher Scientific | Cat# S11223 |
| AF555 streptavidin | Thermo Fisher Scientific | Cat# S21381 |
| AF633 streptavidin | Thermo Fisher Scientific | Cat# S21375 |
| DAPI | Millipore Sigma | Cat# D9542 |
| RNAscope Multiplex Fluorescent v2 Assay | Advanced Cell Diagnostic | Cat# 323110 |
| RNAscope mouse probe mmu-Sst | Advanced Cell Diagnostic | Cat# 461681 |
| RNAscope mouse probe mmu-*Grin3a*-C2 | Advanced Cell Diagnostic | Cat# 1225361-C2 |
| Opal 520 | AKOVA | Cat# OP-001001 |
| Opal 650 | AKOVA | Cat# OP-001006 |
| ProLong Gold Antifade Mountant | Thermo Fisher Scientific | Cat# P10144 |
| RNeasy Plus Micro Kit | Qiagen | Cat# 74034 |
| RNA 6000 Pico kit | Agilent | Cat# 5067-1513 |
| SMARTer Stranded Total RNA-Seq. Kit v2 | Takara Bio | Cat# 634411 |
| Experimental models: Organisms/strains |  |  |
| Mouse: B6;129X1-*Grin3a*<sup>tm1Nnk/J</sup><br>NR3a KO | The Jackson Laboratory<br>[Das1998] | RRID:IMSR_JAX:029974 |
| Mouse: *Grin3a*<sup>fl/fl</sup> | Isabel Perez-Otano |  |
| Mouse: *Grin1*<sup>fl/fl</sup> | The Jackson Laboratory<br>[Tsien1996] | RRID:IMSR_JAX:005246 |
| Mouse: Sst-Cre::*Grin3a*<sup>fl/fl</sup> | Isabel Perez-Otano Lab |  |
| Mouse: Sst<sup>tm2.1(cre)Zjh/J</sup><br>Sst-IRES-Cre | The Jackson Laboratory<br>[Taniguchi2011] | RRID:IMSR_JAX:013044 |
| Mouse: B6.Cg-NPY<sup>tm1(cre)Zman/J</sup><br>NPY-IRES-Cre | The Jackson Laboratory<br>[Milstein2015] | RRID:IMSR_JAX:027851 |
| Mouse: C57BL/6J-Tg(Nkx2-1-cre)2Sand/J<br>Nkx2.1-Cre | The Jackson Laboratory<br>[Xu2008] | RRID:IMSR_JAX:008661 |
| Mouse: Slc32a1<sup>tm2(cre)Lowl/J</sup><br>VGAT-IRES-Cre | The Jackson Laboratory<br>[Vong2011] | RRID:IMSR_JAX:016962 |
| Mouse: B6.Cg-Gt(ROSA)26Sor<sup>tm9(CAG-tdTomato)Hze/J</sup><br>Ai9(RCL-tdT) | The Jackson Laboratory<br>[Madisen2010] | RRID:IMSR_JAX:007909 |
| Mouse: B6.Cg-Gt(ROSA)26Sor<sup>tm14(CAG-tdTomato)Hze/J</sup><br>Ai14(RCL-tdT)-D | The Jackson Laboratory<br>[Madisen2010] | RRID:IMSR_JAX:007914 |
| Mouse: B6.Cg-Gt(ROSA)26Sor<sup>tm40.1(CAG-aop3/EGFP)Hze/J</sup><br>Ai40(RCL-ArchT/EGFP)-D | The Jackson Laboratory | RRID:IMSR_JAX:021188 |
| Publicly available transcriptomic data, integrated and reanalysed |  |  |
| scRNAseq data – processed count file and sample metadata (Allen 2019) | [Hodge2019] | https://transcriptomic-viewer-downloads.s3-us-west-2.amazonaws.com/mouse/transcriptome.zip |
| scRNAseq data – processed count file and sample metadata (Allen 2021) | [Yao2021] | https://assets.nemoarchive.org/dataset/jb2f34y |
| scRNAseq data – processed count file and sample metadata (Harvard 2020) | [Krienen2020] | [GSE151761](https://www.ncbi.nlm.nih.gov/geo/query/acc.cgi?acc=GSE151761) |
| scRNAseq data – processed count file and sample metadata (Karolinska 2018) | [Harris2018] | [GSE99888](https://www.ncbi.nlm.nih.gov/geo/query/acc.cgi?acc=GSE99888) |
| scRNAseq data – processed count file and sample metadata (NIH 2021) | [Mahadevan2021] | [GSE156201](https://www.ncbi.nlm.nih.gov/geo/query/acc.cgi?acc=GSE156201) |
| snRNAseq human data | [Siletti2023] | https://alleninstitute.github.io/abc_atlas_access/descriptions/WHB_dataset.html |
| snRNAseq and MERFISH mouse data | [Yao2023] | https://alleninstitute.github.io/abc_atlas_access/descriptions/WMB_dataset.html |
| Software and algorithms |  |  |
| pClamp 10.6 | Molecular Devices | RRID:SCR_011323 |
| Multiclamp 700B | Molecular Devices | RRID:SCR_018455 |
| FIJI (ImageJ) | NIH | RRID:SCR_002285 |
| Excel | Microsoft | RRID:SCR_016137 |
| ZEN Microscopy Software | Zeiss | RRID:SCR_013672 |
| SPOT 5.2 | SPOT Imaging | RRID:SCR_014313 |
| MATLAB 2023b | MathWorks | RRID:SCR_001622 |
| R v3.5.0 and greater | R Foundation | https://www.R-project.org |
| Seurat v3.4 | [Stuart2019] | https://github.com/satijalab/seurat |
| RStudio IDE | RStudio | https://www.rstudio.com |
| DESeq2 | Bioconductor [Love2014] | RRID:SCR_015687 |

### RESOURCE AVAILABILITY

#### Lead Contact

Further information and requests for resources and reagents should be directed to and will be fulfilled by the lead contact, Kenneth Pelkey.

#### Materials Availability

This study did not generate novel, unique reagents.

#### Data and Code Availability

Data generated during this study are available upon request.

### EXPERIMENTAL MODEL AND STUDY PARTICIPANT DETAILS

#### Mice

All mice experiments were conducted in accordance with animal protocols approved by animal care and use committee (ACUC) at the National Institute of Child Health and Human Development (ASP# 20-045) and at Emory University.

Targeted neuronal subpopulations were achieved by fluorescently labeling, using Cre-recombinase driven expression of floxed reporters. Experimental offspring were maintained as heterozygous crosses from homozygous Cre-recombinase driver Sst-IRES-Cre (RRID:IMSR_JAX:013044)[Taniguchi 2011], NPY-IRES-Cre (RRID:IMSR_JAX:027851)[Milstein 2015], Nkx2.1-Cre (RRID:IMSR_JAX:008661)[Xu 2008], *Grin1* (RRID:IMSR_JAX:005246) [Tsien 1996] mouse lines and homozygous floxed reporters Ai9(RCL-tdT) (RRID:IMSR_JAX:007909)[Madisen 2010], Ai14(RCL-tdT)-D (RRID:IMSR_JAX:007914) [Madisen 2010], Ai40(RCL-ArchT/EGFP)-D (RRID:IMSR_JAX:021188) lines.

#### Non-Human Primates (NHPs)

Adult rhesus macaque and marmoset tissue was obtained as described in [Caccavano 2025]. All experiments were performed in accordance with the ILAR Guide for the Care and Use of Laboratory Animals and were conducted under Animal Study Protocols approved by the ACUC at the National Institute of Mental Health. All procedures adhered to the applicable Federal and local laws, regulations, and standards, including the Animal Welfare Act and Regulations and Public Health Service policy (PHS2002).

### METHOD DETAILS

#### Viral Injections (mouse)

For perinatal injections as described in [Caccavano 2025], P0 mice were anesthetized by hypothermia. Mice were bilaterally injected free-hand with 1 µL of solution containing pAAV(AAV5)-pCAG-FLEX-EGFP-WPRE (Addgene 51502-AAV5, ≥ 7 × 10^12^ vg/mL) in the lateral ventricles: 0.8 mm anterior to lambda and 0.8-1 mm lateral to the sagittal suture, using a 32G microliter Neuros Syringe (Hamilton).

#### Viral Injections and brain harvests in NHPs

As described in [Caccavano 2025], briefly intracranial macaque injections were targeted using stereotaxic coordinates derived from MRI and delivered using a needle guide for enhanced accuracy [Fredericks 2020]. Surgeries were performed under aseptic conditions in a fully equipped operating suite. pAAV-PHP.eB-BiSSTe4-dTomato (Addgene 213944) or pAAV-PHP.eB- BiSSTe10-dTomato (Addgene 213814) (≥ 1×10¹³ vg/mL) was injected into HPC and M1 of 6 rhesus macaques. Within HPC, 30-50 μL of total virus was injected, with 15-25 μL of virus injected at each of 2 locations spaced approximately 2 mm apart in the antero-posterior plane, caudal to the level of the uncus. Within M1, 30-40 μL of total virus was injected, with 10 μL of virus injected at each of 3-4 locations spaced approximately 2 mm apart, targeted via direct visualization.

For NHP brain extraction, (6-8 weeks after virus injection for macaque), animals were sedated with ketamine/midazolam (ketamine 5-15 mg/kg, midazolam 0.05-0.3 mg/kg) and maintained on isoflurane. A deep level of anesthesia was verified by an absence of toe-pinch and corneal reflex response. Prior to brain removal and blocking, NHPs were transcardially perfused with ice-cold sucrose-substituted artificial cerebrospinal fluid (SSaCSF) containing in mM: 90 Sucrose, 80 NaCl, 3.5 KCl, 1.25 NaH_2_PO_4_, 24 NaHCO_3_, 10 Glucose, 0.5 CaCl, 4.5 MgCl_2_ saturated with carbogen (95% O_2_, 5% CO_2_), with osmolarity 310-320 Osm. Skulls were opened and brain was removed and blocked in cold oxygenated SSACSF then sectioned as described below for mouse tissue.

#### Acute brain slice preparation

Postnatal (p5–p10) or juvenile (p11–p21) mice were anesthetized with isoflurane and then decapitated. The brain was dissected in ice-cold high-Mg^2+^ low-Na^+^ ACSF for postnatal mice, sucrose-substituted ACSF for juvenile or adults. The high-Mg^2+^ low-Na^+^ ACSF contains the following (in mM): 130 NaCl, 3.5 KCl, 24 NaHCO_3_, 1.25 NaH_2_PO_4_, 4.5 MgCl, 0.5 CaCl_2_, and 10 glucose with osmolarity 300-310 mOsm. The sucrose-substituted ACSF contains in mM 90 sucrose, 80 NaCl, 3.5 KCl, 24 NaHCO_3_, 1.25 NaH_2_PO_4_, 4.5 MgCl, 0.5 CaCl_2_, and 10 glucose. Horizontal hippocampal slices (300 μm) were sectioned using a VT-1200S vibratome (Leica Microsystems), then transferred and incubated in high-Mg^2+^ low-Na^+^ ACSF at 32°C at least 1 hour before recording. Both ACSFs were saturated with 95% O_2_ and 5% CO_2_.

#### Ex vivo Electrophysiology recording

The brain slices were transferred to an upright microscope (Zeiss Axioskop 2), perfused at 2-3 ml/min with oxygenated ACSF containing in mM 130 NaCl, 3.5 KCl, 24 NaHCO_3_, 1.25 NaH_2_PO_4_, 1.5 MgCl, 2.5 CaCl_2_, and 10 glucose, and maintained at a temperature of 32°C-34°C. Electrodes were pulled from borosilicate glass (World Precision Instruments) to a resistance of 2.5–4 Mohm using a vertical pipette puller (PP-830, Narishige). Whole-cell patch-clamp recordings were made using a Multiclamp 700B amplifier (Molecular Devices), filtered at 3 kHz (Bessel filter), and digitized at 10-25 kHz (Digidata 1550, Molecular Devices) and pClamp 10.6 software; Molecular Devices). Uncompensated series resistance during recordings ranged from 7 to 30 Mohm and was monitored continuously throughout with 5mV voltage steps. In current-clamp mode, cells were hold to −70 mV; in voltage-clamp mode, holding potentials were set to −70 mV for measuring holding current and 0-10 mV for G-PSC and GDP-I. For whole-cell recordings, internal solutions used were as follows. For recording glycine puff response of NGF-INs in [Figure 2C] and action potentials and holding currents in [Figure 3], the internal solution contains the following (in mM): 130 K-gluconate, 5 KCl, 2 MgCl_2_, 2 MgATP, 0.3 NaGTP, 10 HEPES, 0.6 EGTA, and 0.2% biocytin. In other experiments, the internal solution contains in mM 123 CsCH_3_SO_4_, 4.5 NaCl, 4 MgATP, 0.4 NaGTP, 10 HEPES, 1 QX-314, 10 Bapta, and 0.2 % biocytin. The pH of internal solutions was adjusted to 7.4, and osmolarity was adjusted to 285-290 mOsm.

For synaptically isolated conditions, NMDA, AMPA and GABA_A_ receptors blocked by APV 50 μM, DNQX 10 μM, BIC 10 μM and PTX 50 μM respectively. Binding CGP-78608 to the GluN1 subunit does not only inhibit desensitization of eGlyRs, but also inhibits conventional GluN2-containing NMDAR, influencing (excitatory) post-synaptic current. Thus, for recording network activity changes by CGP-78608 application, the brain slices were incubated in 100 μM APV containing ACSF for at least 1 hour prior to and during recording, to exclude influence of conventional NMDAR-mediated current change by CGP on the GDP generation.

NGFCs express Neuro peptide Y (NPY) [Price 2005, Karagiannis 2009, Tricoire 2010], and the combination of the NPY-Cre mouse line and the cell location in the stratum lacunosum moleculare (SLM) enables targeted recording of NGFCs. The labeled NPY-NGF-INs showed late spiking, which is the electrophysiological signature of NGF-INs. For targeting Sst-INs, Sst-IRES-Cre::Ai9-tdTomato or Sst-IRES-Cre::Ai14-tdTomato mice were adopted. In some experiments, putative OLM neurons expressing Sst were targeted based on their morphology such as a horizontally oriented cell body shape, and a deep layer location in the SO near the alveus. Their identification was confirmed by extending the axons to reach the SLM through post hoc cell morphology reconstruction using biocytin staining.

#### Local field potential recording

Local field potential (LFP) recordings were obtained on an interface recording set-up (Scientific System Design Inc.). Brains of adult (>P60) mice were dissected as mentioned above in ice-cold sucrose-substituted ACSF. Horizontal hippocampal slices were sectioned at 350 µm, then transferred to an interface holding chamber with the slices suspended on the surface level of oxygenated 2 mM MgCl/CaCl_2_ ACSF containing the following (in mM): 130 NaCl, 3.5 KCl, 24 NaHCO_3_, 1.25 NaH_2_PO_4_, 2 MgCl, 2 CaCl_2_, and 10 glucose with an osmolarity of 300-310 mOsm. Slices were incubated at 32°C in the holding chamber for 30 minutes, then held at room temperature until recording. Slices were transferred to an upright microscope (Zeiss Stemi 2000- C) and interface chamber, perfused at 2 ml/min with 32°C-34°C oxygenated 2 mM MgCl/CaCl_2_ ACSF containing 50 µM of APV. 3-5 MOhm tip electrodes were filled with 2 mM MgCl/CaCl_2_ ACSF. A Multiclamp 700B amplifier (Molecular Devices) was used to make LFP recordings, filtered at 3 kHz (Bessel filter), and digitized at 10-25 kHz (Axon Digidata 1550, Molecular Devices) to pClamp 10.6 software; Molecular Devices). LFP recordings were conducted in hippocampal CA3 pyramidal cell layer. Spontaneous sharp wave-ripples (SWRs) were recorded in 10-minute epochs for baseline, bath-applied drug (1 µM CGP-78608 or 40 µM EU1180-490), and washout conditions. Recording traces were filtered at 1 Hz (high pass). The frequency of SWRs was determined by the number of SWRs detected during a 2-minute period using Clampfit 10 software (Molecular Devices). Traces were filtered 150-250 Hz to confirm ripples.

#### Post hoc morphological Reconstruction

Slices containing biocytin filled cells were fixed in 4 % paraformaldehyde overnight at 4 °C. Then fixed slices were washed in PBS, permeabilized and incubated with Alexa-488, Alexa-555, or Alexa-633 conjugated streptavidin overnight (Molecular Probes). After incubation, slices were washed multiple times and then were resected to 70μm using a freezing microtome (Microm, Whaltham, MA). Slices were mounted on glass slides (Fisherbrand Superfrost Plus, Fisher Scientific) using Mowiol as a mounting medium. Cells were visualized using epifluorescence microscopy (AX70, Olympus). Confocal images of labelled cells were attained on a Zeiss LSM 780 microscope (NICHD Imaging and Microscopy Core).

#### Immunohistochemistry (IHC)

Mice of different ages (P6, 13, and 6 weeks) were deeply anesthetized with isoflurane and transcardially perfused with 4% paraformaldehyde (PFA) in phosphate buffered saline (PBS) pH 7.4. Brains were removed, post-fixed 1 hour in the same fixative at 4°C and transferred to 30% sucrose in PBS at 4°C for at least 24 hours before sectioning. 30 μm coronal slices were cut and stored at -20°C in cryoprotectant of PBS, 30% ethylene glycol and 30% glycerol in PBS.

Free-floating brain sections were briefly washed in PBS to remove cryoprotectant and then incubated with blocking solution containing 4% normal goat serum (NGS), 1% BSA, 0.1-0.3% Triton X-100 in PBS for 2 hours. Then sections were incubated with primary antibody in 1% NGS, 1% BSA, 0.1-0.3% Triton X-100 in PBS overnight at 4°C. The following primary antibodies were used: rabbit anti-GFP (Synaptic Systems 132002, 1:1000), chicken anti-GFP (Aves labs GFP-1020, 1:1000), rabbit anti-SST (Invitrogen PA585759, 1:500), rabbit anti-nNOS (Abcam ab5586, 1:200).

Sections were then washed extensively with PBS and incubated with fluorophore conjugated secondary antibody in 1% BSA, 0.1% Triton X-100 in PBS for 2-3 hours at room temperature (overnight at 4°C for dendritic analysis). Secondary antibodies used were: anti-rabbit Alexa 488 (Invitrogen A11008 1:750), anti-chicken Alexa 488 (Invitrogen A11039, 1:750) and anti-rabbit Alexa 555 (Invitrogen A21428, 1:500). After PBS washes, sections were counterstained with DAPI, mounted onto Superfrost Plus slides, air-dried, and coverslipped with fluorescence mounting medium.

#### In situ Hybridization (ISH) RNAscope

To prepare snap frozen tissue for ISH, freshly dissected macaque hippocampal tissues were submerged into 2-methylbutane that was pre-chilled on a dry ice/ethanol bath for 5 minutes. Tissue was removed, wrapped in foil, and stored at -80 °C. 10 μm frozen sections were made using a Leica Cryostat, mounted on Superfrost Plus slides (Fisher Scientific Cat#12-550-15) and stored at -80 °C.

Target probes were designed and manufactured by Advanced Cell Diagnostic (ACD): mmu- SST(Cat#461681), mmu-G*rin3a*-C2(Cat#1225361-C2). ISH was performed following the RNAscope Multiplex Fluorescent v2 Assay instructions provided by ACD. Briefly, frozen thin sections were post-fixed in 4% PFA and dehydrated sequentially in 50%, 70% and 100% ethanol. After H2O2 and Protease IV treatment, sections were incubated with probes at 40 °C for 2 hours. Probed signals were detected using RNAscope Multiplex Detection v2 kit (ACD, Cat# 323110). Opal 520 (AKOYA, Cat# OP-001001) and Opal 650 (AKOYA, Cat# OP-001006) were used to detect C1 (sst) and C2 (*Grin3a*) probes, respectively. After DAPI staining, sections were covered using ProLong Gold Antifade Mountant (ThermoFisher Scientific, Cat# P10144) and cured in darkness before imaging.

RNAscope images were acquired on a SLIDEVIEW VS200 slide scanner (Olympus) mounted with an ORCA-Fusion Digital Camera (HAMAMATSU). Hippocampal regions were scanned at 20x and 40x for counting and signal colocalization analysis.

#### RiboTag

This assay was performed as previously described[Sanz 2009, Mahadevan 2020, Mahadevan 2021]. RNA bound with anti-HA immunoprecipitates and RNA from bulk tissue were purified using RNeasy Plus Micro Kit (Qiagen, Cat# 74034) and the quality of RNA was measured using RNA 6000 Pico kit (Agilent, Cat# 5067-1513) and 2100 Bioanalyzer system (Agilent, Cat# G2939BA). cDNA libraries were constructed from 250 pg RNA using the SMARTer Stranded Total RNA-Seq. Kit v2 (Takara Bio, Cat# 634411) from samples with RNA Integrity Numbers > 6.

Sequencing of the libraries were performed on the Illumina HiSeq 2500, at 50 million 2 × 100 bp paired-end reads per sample. ∼75% of reads were uniquely mapped to genomic features in the reference genome. Bioconductor package DESeq2[Love 2014] was used to identify differentially expressed genes (DEG). This package allows for statistical determination of DEGs using a negative binomial distribution model. The resulting values were then adjusted using the Benjamini and Hochberg’s method for controlling the false discovery rate.

#### Quantification and statistical analysis

Data analysis and visualization was conducted by homemade MATLAB codes. GDP-I was detected using simple threshold method after gaussian filtering, and G-PSC detection was achieved by template event detection in Clampfit 10.6. Paired t-test was selected for change of activity by chemical or light stimuli, and Wilcoxon rank sum test was used for comparison between different conditions (*: 0.01 to 0.05, **: 0.001 to 0.01, ***: < 0.001). Summary data are presented as mean ± SEM.

#### Multi-institute single-cell transcriptome analyses of GABAergic interneurons

Single-nucleus (sn) / single-cell (sc) transcriptomes from mouse cortical and hippocampal regions were downloaded from studies spanning research institutes (Allen Institute: [Hodge 2019, Yao 2021], Harvard Medical School: [Krienen 2020], Karolinska Institute: [Harris 2018], NIH: [Mahadevan 2021]). HDF5*/*raw files from these sources were downloaded and converted into Seurat v3-compatible format using custom scripts in R package as described earlier [Chittajallu 2020, Ekins 2020]. Basic integration, processing and visualization of the scRNA-seq data were performed with the Seurat v3 [Butler 2018, Stuart 2019] in R environment using R Studio. Briefly, data was preprocessed by removing cells with less than 200 detectable genes and with reads mapping to mitochondrial genes comprising more than 0.05% of total reads.

Gene expression counts were then log normalized and the resulting expression matrix was scaled to 10,000 transcripts per cell. Next, the expression values of each gene across all the cells (z-score transformation) were utilized for dimensionality reduction. These were performed using the NormalizeData() and ScaleData() functions in Seurat with default parameters. Next, we identified the top 2000 genes that exhibited high variability across cells and therefore represent unique features of the cell clusters, for downstream analysis using the FindVariableGenes() function, by applying selection.method = “vst” feature. Next, principal component analysis (PCA) was performed, and the top 20 principal components were stored.

We clustered the cells using the FindClusters() with a clustering resolution set to 2.0. Using the FindAllMarkers() function at cutoff of genes expressed in 25% of cells, at a logfc.threshold = 0.25, we identified the genes that are expressed in each cluster. For each cell type we used the well-established marker genes that uniquely established the cell identity, as previously described in the respective studies. Subsetting only the Gad1 clusters expressing a combination of Lhx6, Pvalb, Sst, Id2 were annotated as MGE-derived GABAergic interneurons, and a combination of Lhx6-negative, Sncg, Vip, Calb2, Meis2, Cnr1, Ntng1, Ndnf were annotated as CGE-derived GABAergic interneurons. The Neurogliaform interneurons that are Lhx6^+^/Lamp5^+^/Id2^+^ are annotated as MGE-derived (NGFC.M); and those that are Lhx6^-^/Lamp5^+^/Id2^+^/Ndnf^+^ are annotated as CGE-derived (NGFC.C). In total we used 86,852 GABA- subset sc-transcriptomes in this study. Expression levels of various GABAergic inteneuron positive control genes and expression of *Grin3a* across the GABAergic interneuron clusters are plotted using DotPlot function in Seurat [Figure 2A].

#### Analyses of publicly available transcriptomic data

For cross-species comparison of *GRIN1* and *GRIN3A* expression, we utilized a publicly available single-nucleus RNA sequencing (snRNA-seq) dataset comparing across neocortical INs in mouse, marmoset, macaque, and human [Krienen 2020]. We probed the mean expression across four neocortical IN subpopulations (PV, SST, ID2, VIP) with a gene panel including *GRIN1*, *GRIN3A*, and distinguishing markers *PVALB*, *SST*, *LAMP5*, and *VIP*.

For further analysis into human single cell expression, we utilized the Allen Institute whole human brain dataset of the single-cell transcriptomes of ∼3.4 million cells across the entire human brain [Siletti 2023]. To probe our genes and regions of interest, we adapted Python notebooks provided for accessing the Allen Brain Cell (ABC) Atlas (https://alleninstitute.github.io/abc_atlas_access/intro.html). The log2-normalized expression matrices were downloaded and filtered through several criteria to reach the cell types of interest. First, only neurons were considered (∼2.5m cells), which were limited to the superclusters “MGE interneuron”, “CGE interneuron”, “LAMP5-LHX6 and Chandelier”, “Hippocampal CA1-3”, “Hippocampal CA4”, and “Hippocampal dentate gyrus” (∼648k cells), and finally to the hippocampal anatomical division (∼204k cells). Analysis of PNs was limited to the supercluster level, while INs were further analyzed down to the cluster level to meaningfully group clusters into identifiable interneuron subpopulations (e.g. PV, SST, LAMP5). We calculated the mean log2-normalized expression values across clusters within this subset of cells for a gene panel including: *GAD1*, *GAD2*, *SLC17A7* (VGluT1), *SLC17A6* (VGluT2), *LHX6*, *PROX1*, *PVALB*, *SST*, *VIP*, *LAMP5*, *CCK*, *SNCG*. Using a mean threshold = 2, MGE-PV clusters were defined as those expressing *LHX6*+, *PVALB*+ (MGE_261/263, LLC_264/266, 1480 cells), MGE-SST clusters as *LHX6*+, *SST*+ (MGE_263/239-254/258, LLC_271, 12935 cells), MGE-LAMP5 as *LHX6*+, *LAMP5*+ (LLC_266-275, 7663 cells), CGE-LAMP as CGE *LAMP5*+ (CGE_286-288, 6497 cells), CGE-SNCG as CGE *SNCG*+ (CGE_276/277/279/281/285/291, 6604 cells), and CGE-VIP as CGE *VIP*+ (CGE_276/280/281/284/285/289-296, 16007 cells). We next downloaded the single-cell gene expression across these defined groups for a gene panel including *GRIN1* and *GRIN3A* to create violin and dot plots via Graphpad Prism. All adapted Python notebooks are available on Github (https://github.com/acaccavano/ABC_Atlas_Kim2025).

---

**S1.**
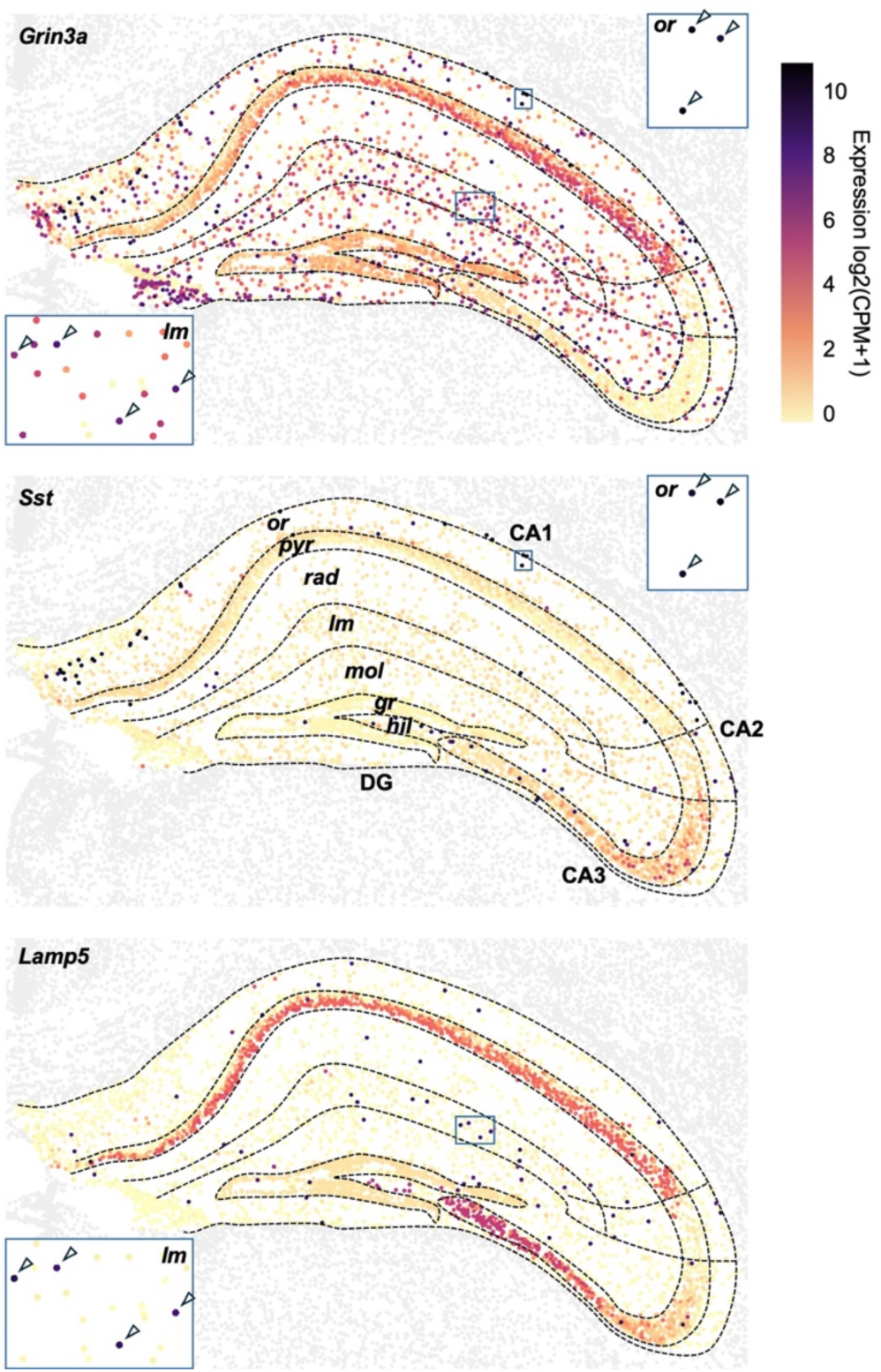
***Grin3a* is absent in CA3 PNs and enriched in *Sst* and *Lamp5* expressing cells.** Publicly available spatial transcriptomics MERFISH data from the Allen Brain Cell Atlas with imputed gene expression [Yao 2023]. Visualized gene expression restricted to the hippocampal formation, showing (top) *Grin3a*, (middle) *Sst*, and (bottom) *Lamp5*. Insets highlight expression of putative Sst-INs in stratum oriens (or) and Lamp5-INs in stratum lacunosum-moleculare (lm). Arrows denote significant expression of Grin3A transcript in SST or Lamp5 expressing INs.

**S2.**
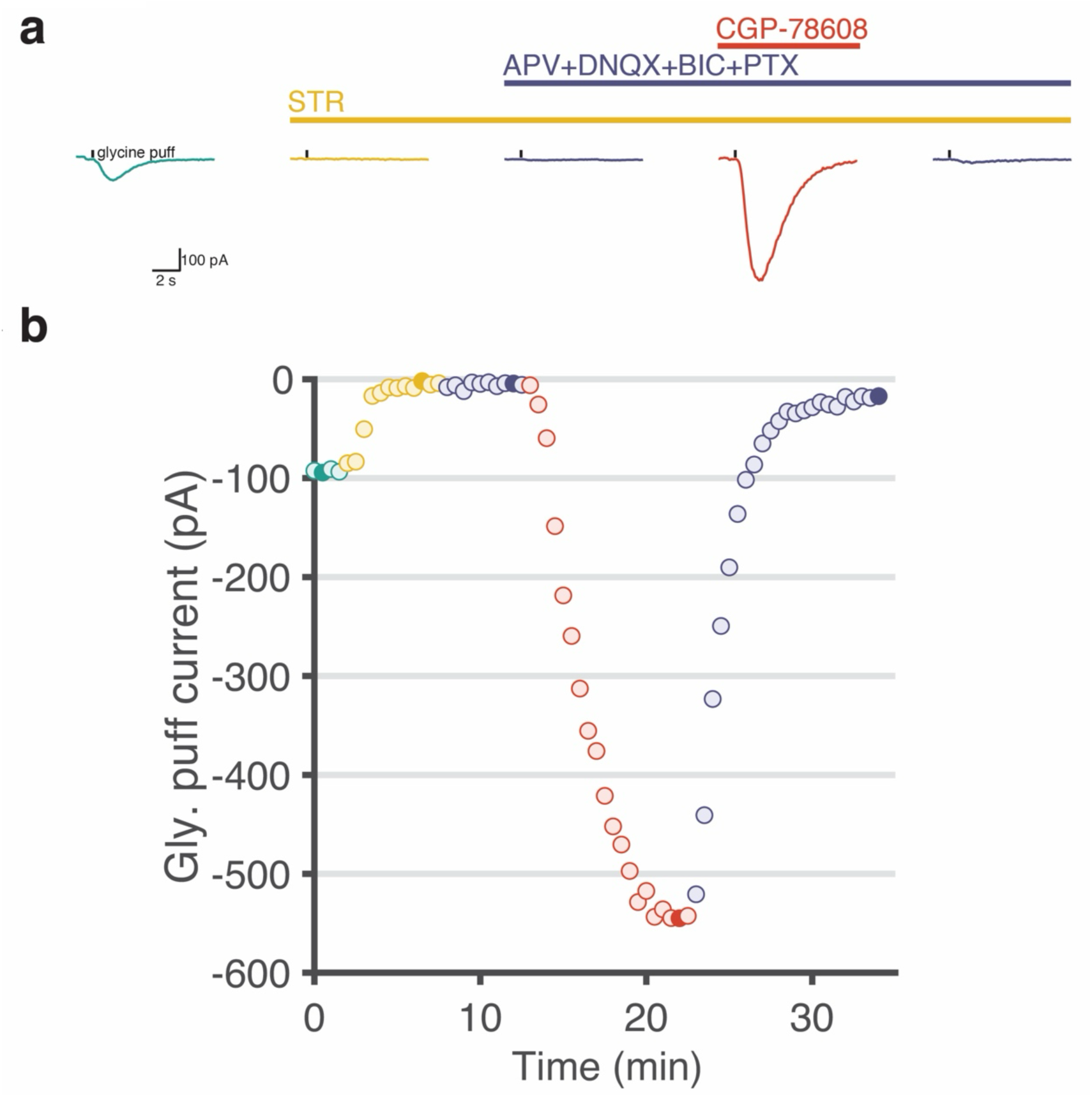
Experimental procedure employed to isolate strychnine-insensitive glycine evoked eGlyR mediated depolarizing currents. (a) Example current traces from a CA1 Sst-IN with glycine puff responses in presence of blockers and CGP-78608, filled circles in (b). Strychnine sensitive eGlyR-mediated current prominently contributed the glycine response current in ACSF. (b) Glycine puff currents in antagonists and CGP-78608. * STR 1 μM, APV 50 μM, DNQX 10 μM, BIC 10 uM and PTX 50 μM.

**S3.**
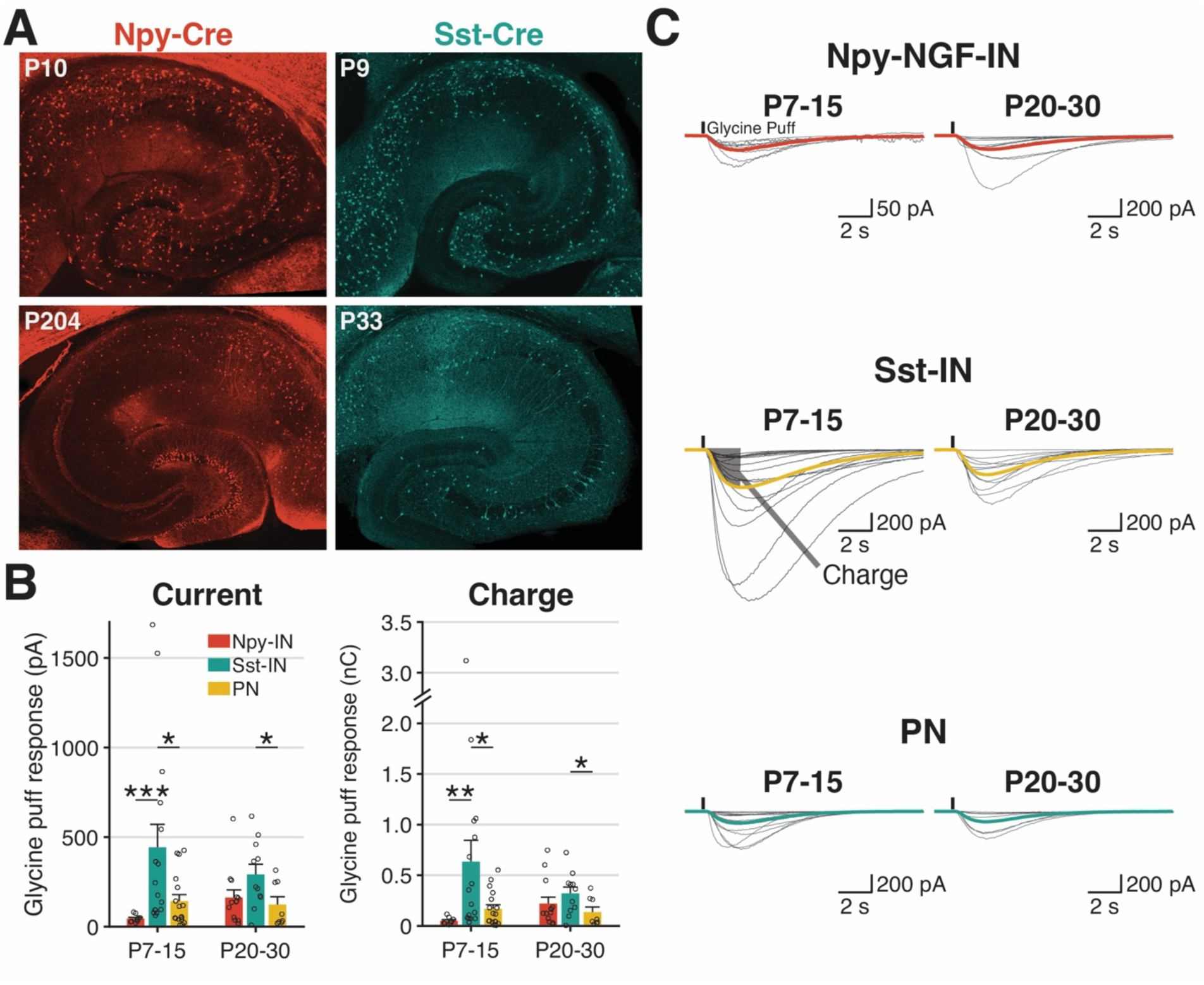
eGlyRs mediate functional responses in SST and NPY-NGFC INs throughout development. **A.** Sample images of postnatal (top)/adulthood (bottom), NPY-IRES-Cre (left)/Sst-IRES-Cre (right) mouse hippocampal slices. **B.** (left) Glycine puff response currents in NPY-INs, Sst-INs, and PNs from postnatal and adulthood mice. (right) Glycine puff response charge transfer in NPY-INs, Sst- INs, and PNs from postnatal and adulthood mice. **C.** Glycine puff responses in NPY -INs, Sst-INs and PNs in different ages. Thick lines are average puff responses of cells and gray lines are average puff responses of trials in each cell.

**S4.**
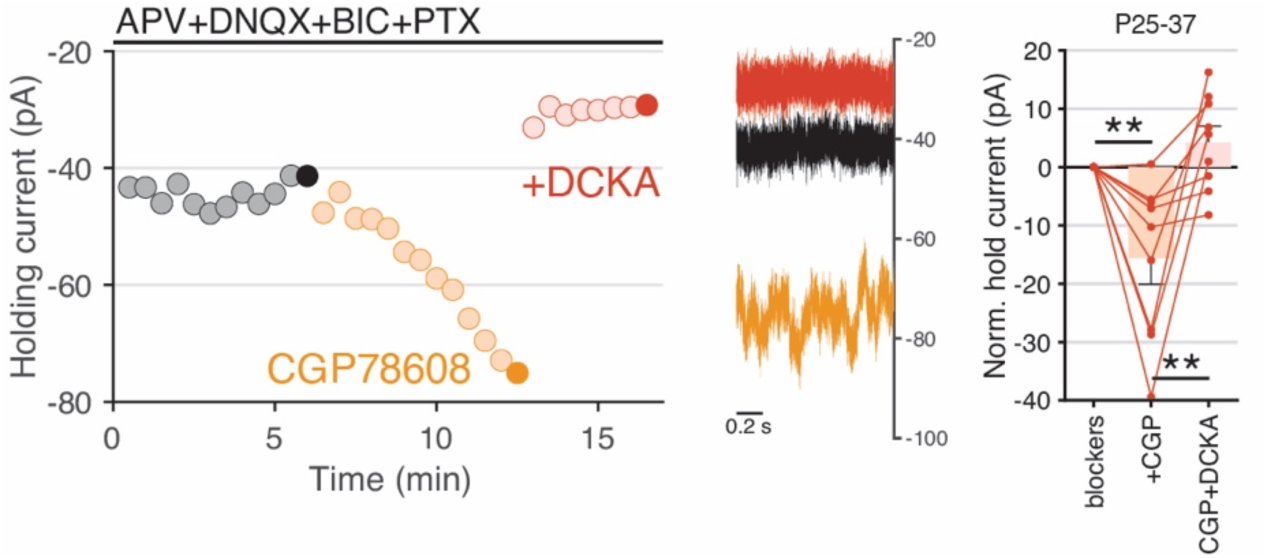
DCKA sensitivity of CGP78608-induced change holding current in hippocampal interneurons. Shown are the time course (left) and traces (middle) from a representative sample NGFC recording illustrating increased holding current (V_h_=-70mV) in response to CGP78608 and subsequent reduction of holding current upon DCKA application. At right, is the group data summarizing the effects of CGP and subsequent DCKA application upon holding currents for Sst- INs and NGFCs combined.

**S5.**
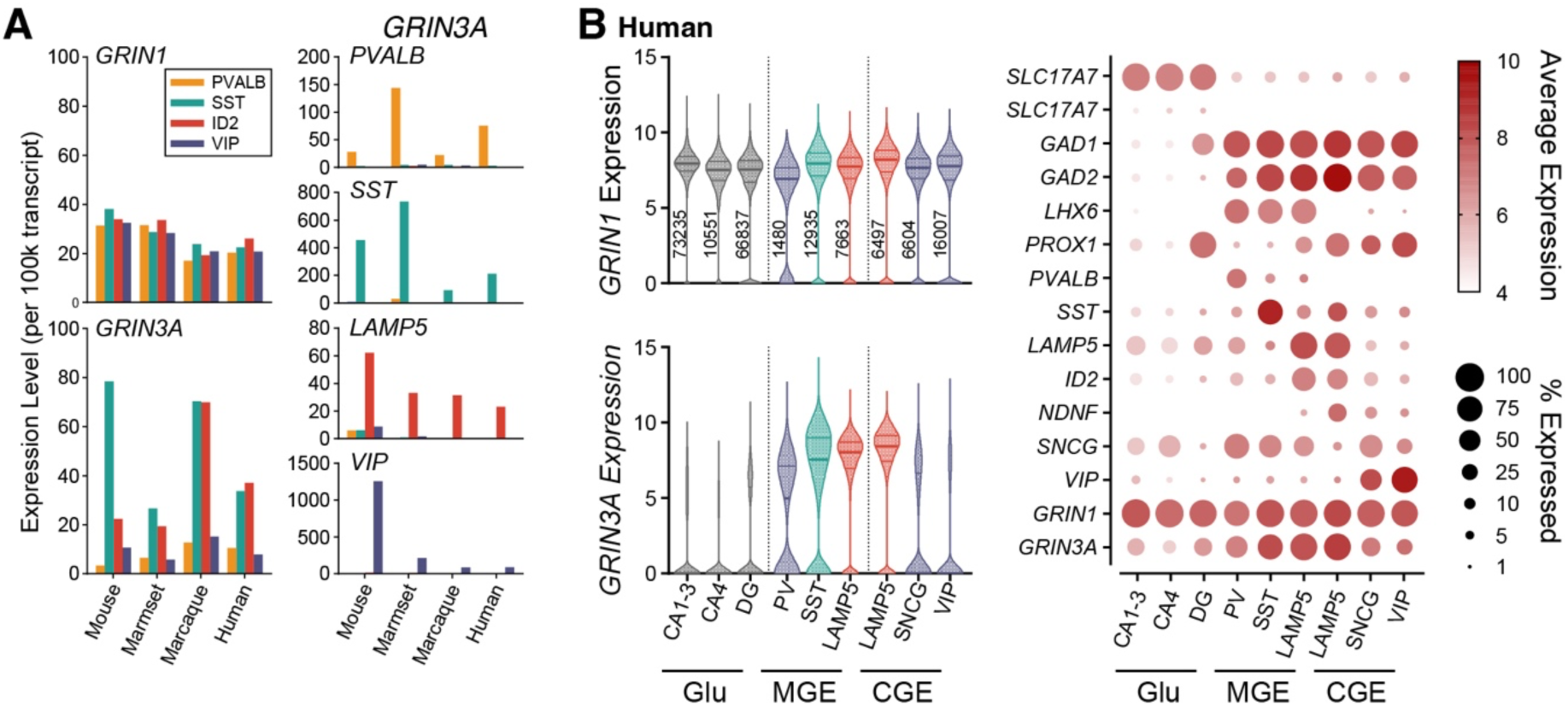
***GRIN1* and *GRIN3A* expression cross species. A.** Publicly-available snRNA-seq mean expression in neocortical IN subpopulations PVALB (orange), SST (cyan), ID2/LAMP5 (red) and VIP (purple) across mouse, marmoset, macaque, and human [Krienen 2020], for (left) *GRIN1* and *GRIN3A* and (right) distinguishing markers *PVALB*, *SST*, *LAMP5*, and *VIP*. **B.** Publicly-available snRNA-seq of human hippocampal neurons [Siletti 2023], with (left) *GRIN1* and *GRIN3A* expression in glutamatergic (Glu) superclusters and grouped MGE and CGE clusters (see Methods), and (right) dot plot of distinguishing marker genes and *GRIN1*, *GRIN3A*.

